# Structure-based inhibitors reveal roles for the clathrin terminal domain and its W-box binding site in CME

**DOI:** 10.1101/2020.01.24.919092

**Authors:** Zhiming Chen, Rosa Mino, Marcel Mettlen, Peter Michaely, Madhura Bhave, Dana Kim Reed, Sandra L. Schmid

## Abstract

Clathrin-mediated endocytosis (CME) occurs via the formation of clathrin-coated vesicles from clathrin-coated pits (CCPs). Clathrin is recruited to CCPs through interactions between the AP2 complex and its N-terminal domain (TD), which in turn recruits endocytic accessory proteins. Inhibitors of CME that interfere with clathrin function have been described, but their specificity and mechanisms of action are unclear. Here we show that overexpression of the TD with or without the distal leg specifically inhibits CME and CCP dynamics by perturbing clathrin interactions with AP2 and SNX9. We designed small membrane-penetrating peptides that mimic the four known binding sites on the TD. A peptide, Wbox2, designed to mimic to the W-box motif binding surface on TD binds to SNX9 and AP2, and potently and acutely inhibits CME, while not perturbing AP1-dependent lysosomal trafficking from the Golgi or bulk, fluid phase endocytosis.

**Summary:** Chen et al define the role the N-terminal domain (TD) of clathrin heavy chain in early and late stages of clathrin-mediated endocytosis, and guided by its structure, design a membrane-penetrating peptide, Wbox2, that acutely and potently inhibits CME.

## Introduction

Clathrin-mediated endocytosis (CME) is the predominant route of receptor entry into cells (Mettlen et al., 2018; Schmid and McMahon, 2007). Clathrin triskelia and AP2 complexes are key constituents of the assembled clathrin-coated pits (CCPs) (Brodsky et al., 2001). The AP2 complexes are multifunctional heterotetramers that: 1) recruit and trigger the assembly of clathrin on the plasma membrane (Cocucci et al., 2012; Edeling et al., 2006; Godlee and Kaksonen, 2013; Kelly et al., 2014; Owen et al., 2000; Shih et al., 1995), 2) recognize cargo receptors, e.g. transferrin receptor (TfnR) (Kelly et al., 2008; Mattera et al., 2011; Ohno et al., 1996; Owen and Evans, 1998; Traub and Bonifacino, 2013), and 3) recruit a myriad of endocytic accessory proteins (EAPs) (Merrifield and Kaksonen, 2014; Owen et al., 1999; Praefcke et al., 2004; Schmid et al., 2006; Traub et al., 1999). Clathrin triskelions bear three clathrin heavy chains (CHCs), each of which contain a proximal leg that binds clathrin light chains (CLCs), a distal leg, and an N-terminal domain (TD) that binds AP2 and a subset of EAPs (Fig. 1A) (Kirchhausen and Harrison, 1981; Royle, 2006; Ungewickell and Branton, 1981). Two antiparallel proximal and two antiparallel distal legs form a polygonal edge of the clathrin lattice and provide rigidity to the coat (Musacchio et al., 1999). The TD is a β-propeller comprised of seven WD40 repeats that generate binding sites for multiple protein interactions identified *in vitro* (Dell’Angelica, 2001; Lemmon and Traub, 2012). However, TD mutations that perturb these interaction surfaces do not inhibit CME, raising doubts as to their cellular functions (Collette et al., 2009; von Kleist et al., 2011; Willox and Royle, 2012). Further studies are needed to understand the role of the TD in CME.

**Figure 1.**
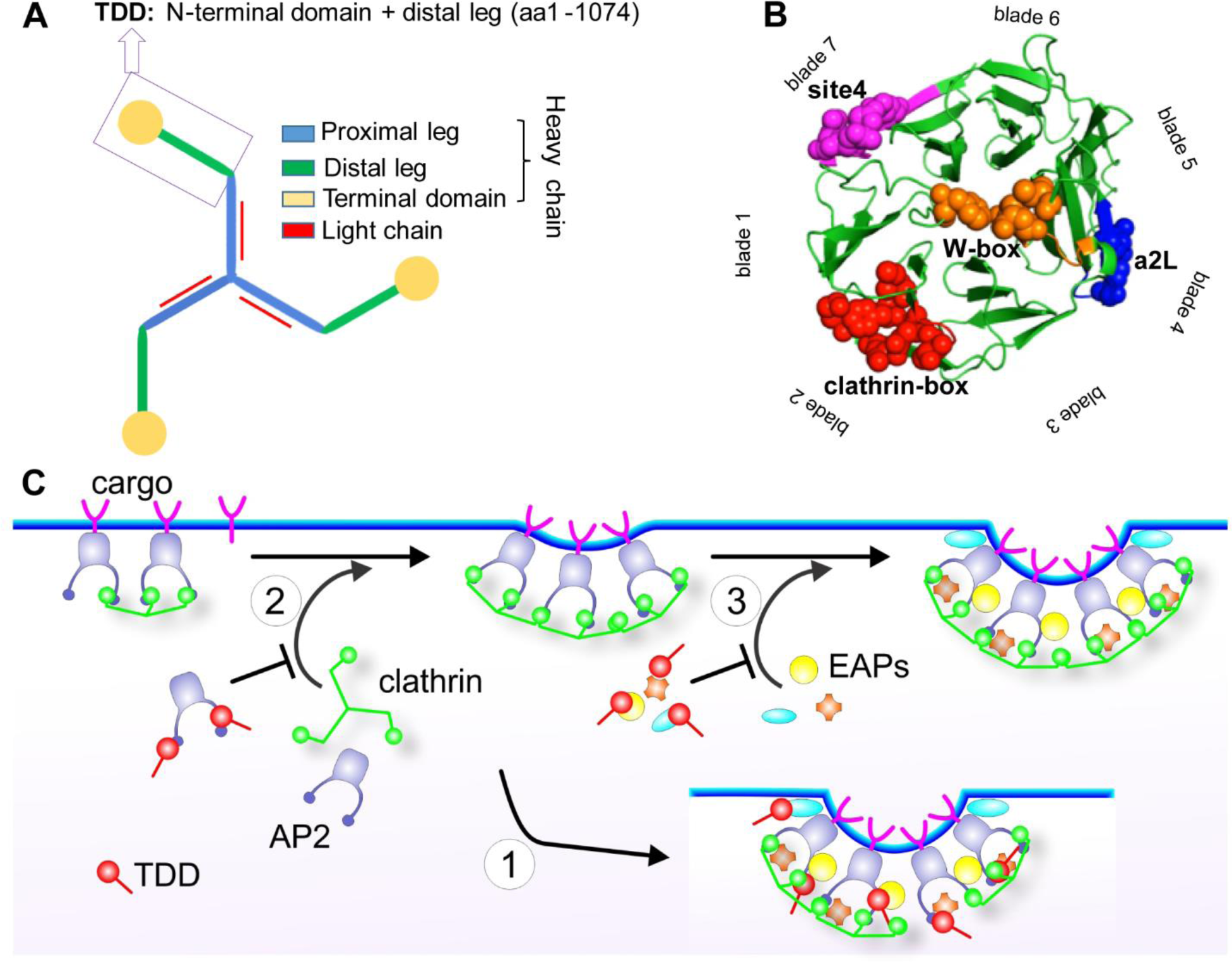
TDD structure and three possible mechanisms of CME inhibition. (A) Domain structure of clathrin triskelion. Box indicates the N-terminal domain and distal leg of clathrin heavy chain (TDD) construct used in this study. (B) Terminal domain of clathrin heavy chain (PDB: 1BPO). Four reported binding sites are labeled with different colors and key functional residues are shown as spheres. (C) Cartoon to illustrate potential TDD inhibitory mechanisms: (1) TDD is incorporated into and destabilizes/weakens the clathrin coat, thus inhibiting CCP maturation; (2) TDD competes for AP2 and inhibits AP2-clathrin interactions; (3) TDD competes for other EAPs required for CCP growth and maturation.

Interfering with clathrin function can suppress CME. Initial studies involved overexpression of the ‘clathrin hub’ that retains only proximal legs. The hub inhibits CME by sequestering clathrin light chains (Bennett et al., 2001; Liu et al., 1998); however, inhibition requires high levels of expression (∼15-fold) and long incubation times (>20 hrs) to allow for turnover of endogenous CHCs and sequestration of newly synthesized CLCs. Moreover, hub overexpression causes a dramatic redistribution of endosomes and blocks cargo transport from the Golgi (Bennett et al., 2001; Liu et al., 1998). A second approach utilizes small molecule inhibitors called ‘Pitstops’, which are reported to block the clathrin-box motif binding site on TD (von Kleist et al., 2011). Pitstops were identified in a screen for chemical inhibitors of TD-EAP interactions; however, subsequent studies have questioned the potency and specificity of Pitstops in clathrin targeting and CME inhibition (Lemmon and Traub, 2012; Liashkovich et al., 2015; Smith et al., 2013; Willox et al., 2014). In addition to the controversy as to the mechanism of action, Pitstop is cytotoxic at concentrations not much higher than those required to inhibit CME (Rosselli-Murai et al., 2018). Thus, there remains a need for specific inhibitors of CME that work by rapid, well-defined mechanisms.

Recent genetic studies have revealed the existence of de novo frame shift mutations resulting in the expression of C-terminally truncated CHC and linked to epilepsy, neurodevelopmental defects and intellectually disabilities (DeMari et al., 2016; Hamdan et al., 2017). Here, we show that overexpression of N-terminal fragments of clathrin heavy chain (TD ± distal leg) corresponding to these frame shift mutations, specifically inhibit CME through interference with AP2 and SNX9 interactions and perturb both early and late stages of CCP maturation.

To identify critical surfaces on TD required for these effects on CME, we designed small membrane-penetrating peptide mimics of the four binding sites on TD and show that a peptide corresponding to the SNX9-binding W-box motif binding site binds directly to SNX9 and potently inhibits CME. Unexpectedly this peptide, named Wbox2, also binds AP2 and inhibits AP2-clathrin interactions; thus phenocopying the effects of TDD overexpression on both early and late stages of CCP maturation.

## Results

To develop a potent inhibitor for CME, we thought to destabilize clathrin lattices or to inhibit clathrin interactions with AP2 or EAPs. The TDD fragment (Fig. 1A) was chosen as a starting point because this region of CHC encompasses both the TD, which is responsible for interactions with AP2 and EAPs (Lemmon and Traub, 2012; Willox and Royle, 2012) (Fig. 1B), and the distal leg regions of CHC involved in clathrin self-assembly (Musacchio et al., 1999). Moreover, recent genetic studies have revealed numerous frameshift mutations in the gene encoding CHC linked to neurological diseases that result in the expression of N-terminal fragments corresponding to our TDD construct. We reasoned that TDD could inhibit CME by three distinct, but not mutually exclusive, mechanisms. First, because the distal leg is a key constituent of polygonal edges of the clathrin lattice that contribute to the rigidity of the assembled coat (Greene et al., 2000; Musacchio et al., 1999), overexpressed TDD could be incorporated into and destabilize clathrin-coated structures by displacing intact clathrin triskelia (Fig. 1C, Mechanism 1). Second, TDD could compete for AP2 binding and inhibit clathrin recruitment and assembly (Fig. 1C, Mechanism 2). Lastly, TDD overexpression could interfere with interactions between clathrin and other EAPs, thereby inhibiting CCP nucleation, stabilization and maturation (Fig. 1C, Mechanism 3). The TDD fragment was cloned into a tetracycline (Tet) regulated (Tet-off) adenoviral system and cells were co-infected with adenovirus encoding the Tet-regulatable transcription factor, tTA. The amounts of TDD and helper tTA adeno-viruses used were optimized for uniform infection of ARPE cells (Fig. S1A-B), such that the levels of TDD expression could be regulated by adjusting Tet concentration (Fig. 2A).

**Figure 2.**
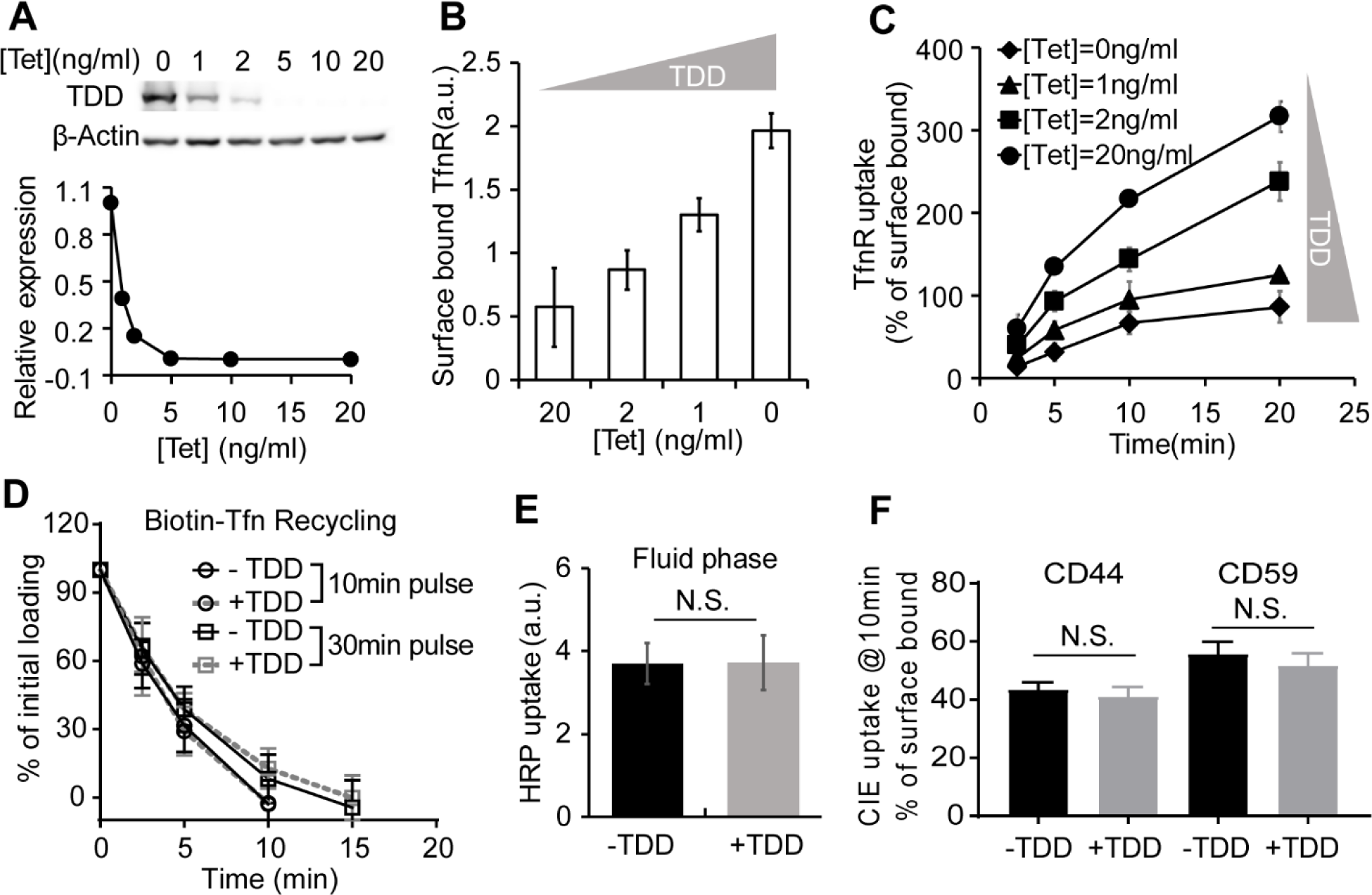
TDD specifically inhibits CME in a concentration-dependent manner. (A) Regulation of TDD expression was achieved by adjusting the tetracycline (Tet) concentration in the Tet-off system. 350,000 ARPE cells in a 6 well plate were infected with 12μl TDD recombinant adenovirus and 12μl tTA recombinant adenoviruses in the presence of the indicated concentrations of Tet. (B-C) Effects of increasing levels of TDD expression on (B) the levels of surface bound TfnR and (C) the rate and efficiency of transferrin receptor (TfnR) uptake assayed in adenovirus-infected ARPE cells in the presence of the indicated concentrations of Tet. (D) Effects of TDD expression (0ng/ml Tet) on Biotin-Transferrin recycling of either a 10 min and 30min pulse in adenovirus-infected ARPE/HPV cells with or without Tet. (E-F) Effect of TDD expression (0ng/ml Tet) on (E) the fluid phase uptake of HRP or (F) clathrin-independent endocytosis of CD44 or CD59 in ARPE/HPV cells. Data ± standard deviations are from N=4 replicates.

### TDD specifically inhibits CME in a concentration-dependent manner

Increasing TDD expression inhibited CME as evidenced by increased surface levels of TfnR and surface-bound transferrin (Tfn) and by decreased TfnR/Tfn internalization (Fig. 2B-C, S1C-F). At maximum TDD expression, the endocytic efficiency of TfnR was inhibited by 80% (Fig. 2C). Recycling of Tfn was not significantly perturbed by TDD expression (Fig. 2D). Similarly, clathrin independent endocytic pathways were not significantly inhibited, as measured using either HRP as a marker for fluid phase uptake, or CD44 and CD59 as markers for clathrin-independent endocytosis (CIE) (Fig. 2E-F). Thus, overexpressed TDD functioned as a potent and specific inhibitor of CME.

### TDD inhibits CCP stabilization and maturation

To define which stage(s) of CME are perturbed by TDD overexpression, we used quantitative live-cell Total Internal Reflection Fluorescence Microscopy (TIRFM) to visualize and analyze CCP dynamics in ARPE/HPV cells stably expressing eGFP-tagged clathrin light chain a (eGFP-CLCa) (Mettlen and Danuser, 2014). TDD overexpression sharply suppressed CCP dynamics. The kymographs of time-lapse imaging showed that large, static clathrin structures predominated when TDD was overexpressed (Fig. 3A-B, Videos S1-2), indicative of a late block in CCP maturation. FRAP analysis revealed that clathrin turnover in these static structures was also impaired (Fig. S2A-C). We also noted a dynamic subpopulation of CCPs (arrowheads, Fig. 3B), which although visibly obscured by the large static CCPs in kymographs, nonetheless represent the majority of total clathrin-labeled structures detected. Thus, we investigated the effects of TDD on earlier stages of CME using cmeAnalysis (Aguet et al., 2013; Jaqaman et al., 2008; Loerke et al., 2011) to quantify the dynamic behaviors of this subpopulation. When TDD was overexpressed, CCP initiation rates dramatically decreased with a corresponding increase in dim and transient clathrin-coated structures (i.e. subthreshold CCSs) (Fig. 3C). The rate and extent of clathrin recruitment (Fig. 3D-E) also decreased. Together, these changes led to a shift toward dimmer structures (Fig. 3F). Finally, the remaining dynamic CCPs also exhibited increased lifetimes upon TDD overexpression (Fig. 3G-H). Thus, we conclude that TDD overexpression inhibits early stages of CCP assembly and stabilization, as well as late stages of CCP maturation.

**Figure 3.**
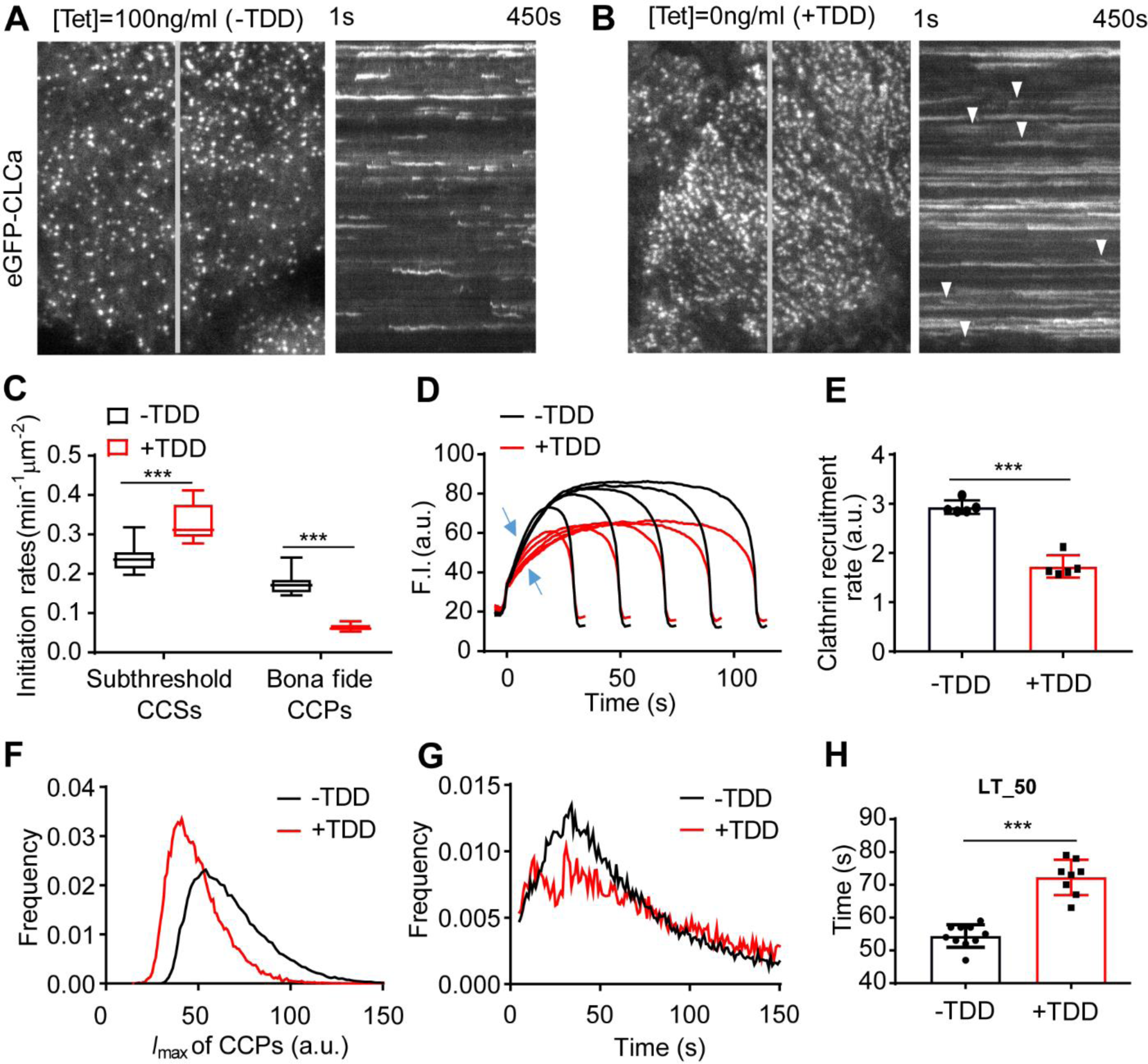
Effects of TDD expression on CCP dynamics. (A-B) Single frame from TIRFM movies (7.5min/movie, see Videos S1-2) and corresponding kymographs from region indicated by gray line of adenovirally-infected ARPE/HPV cells expressing eGFP-tagged clathrin light chain a (eGFP-CLCa) without (A) and with (B) TDD expression. Arrowheads point to examples of dynamic CCSs subjected to cmeAnalysis. (C) Effect of TDD expression on the growth and stabilization of CCPs, as measured by (C) the rates of initiation of subthreshold clathrin-coated structures (CCSs) and *bona fide* CCPs. N = 10 movies for student’s *t-test*. (D) Mean clathrin fluorescence intensity traces in lifetime cohorts of analyzed clathrin structures and (E) corresponding rate of clathrin recruitment determined by the mean slopes of growth phase (from t = 3s to t = 8s) indicated by blue arrows in (D). Each dot represents the mean slope of a cohort. (F) Maximum fluorescence intensity (*I*max) distribution of analyzed CCPs. (G) Lifetime distribution of *bona fide* CCPs. (H) Median lifetime (LT_50) of analyzed CCPs. Each dot represents a movie. Number of dynamic tracks analyzed: 124453 for –TDD and 99846 for +TDD. Error bars are standard deviations.

### TDD is not incorporated into clathrin/AP2 coats but binds to AP2 and SNX9

We next probed the mechanism(s) by which TDD inhibited CME (Fig. 1C). TDD overexpression did not alter endogenous expressions of clathrin and AP2 (Fig. S3A). Unlike the clathrin hub (Bennett et al., 2001; Liu et al., 1998), TDD was recruited to cell membranes (Fig. 4A); however, subcellular fractionation (Fig. 4B, Fig. S3B-C) and immunofluorescence (Fig. S3D) established that TDD was not incorporated into clathrin/AP2 coats. These observations ruled out the possibility that TDD might destabilize clathrin coats (Mechanism 1, Fig 1C), leaving the possibility that TDD might compete with clathrin for binding to AP2 (Mechanism 2) and/or EAPs (Mechanism 3). In support of an AP2/EAP binding mechanism, the TD alone was sufficient to suppress CCP initiation (Fig. 4C).

**Figure 4.**
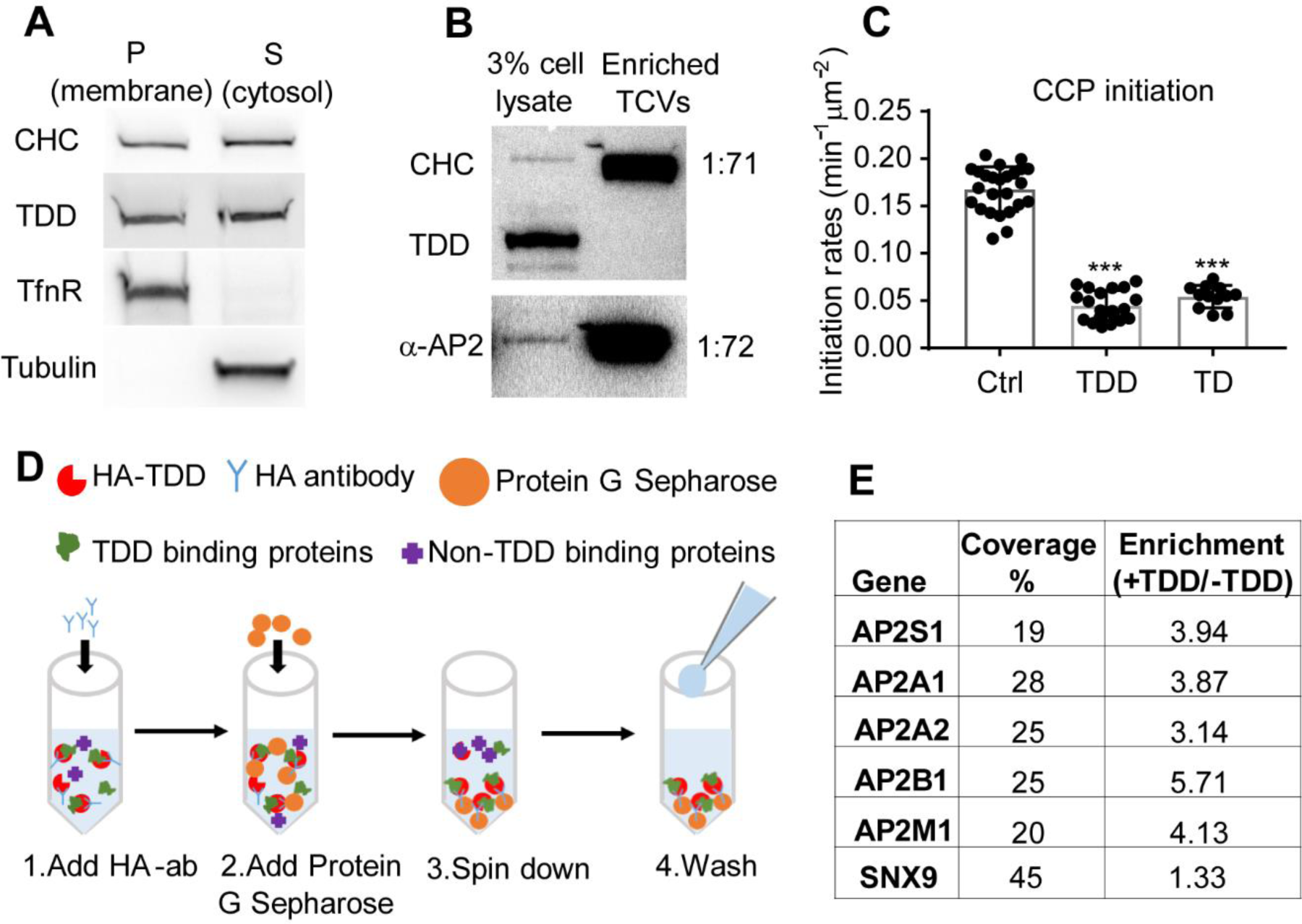
Defining the mechanism of TDD inhibition. (A) Subcellular fractionation of ARPE cells expressing TDD into membrane and cytosolic fractions. (B) Subcellular fractions to enrich for TX-100-resistant clathrin/AP2 coats (TCVs) from of ARPE/HPV cells expressing TDD. (C) TIRFM imaging and quantification of the clathrin dynamics in ARPE/HPV eGFP-CLCa cells indicates that TD phenocopies TDD by reducing the initiation density of *bona fide* CCPs to the same extent. (D) Cartoon showing the procedure used for immunoprecipitation of HA-TDD from ARPE cell lysate (with or without HA-TDD expression) using HA antibody and Protein G Sepharose 4 Fast Flow antibody purification resin. MassSpec analysis was applied to the co-IP samples for protein ID detection. (E) AP2 and SNX9 were identified as the only CME related protein targets that were reproducibly enriched in the TDD co-IPs. Values are average of three independent trials.

We next performed co-immunoprecipitation (co-IP) of HA-tagged TDD from cell lysates (Fig. 4D) to identify *in vivo* binding partners. Three independent trials identified SNX9 and the AP2 complex as the principal CME-related proteins that were reproducibly enriched in TDD vs control co-IPs (Fig. 4E, Table S1). All four subunits of AP2 including both isoforms of the α subunit were identified with high confidence. Although the degree of enrichment of SNX9 was comparatively low, SNX9 was robustly pulled down based on high sequence coverage (Fig. 4E). Surprisingly, the AP1 subunits were either not detected or at very low abundance as compared to AP2 subunits (Table S2). Indeed, TDD overexpression did not alter AP1 localization (Fig. S3E) or Golgi structure (Fig. S3F-G). While the pull-down experiments may have missed weak binding partners, the data suggest that TDD inhibits CME through interference of clathrin interactions with AP2 and SNX9.

### TDD inhibits clathrin-AP2 interactions

To test the effects of TDD overexpression on AP2 function, we tracked the dynamics of AP2 structures by TIRFM in ARPE/HPV cells expressing α-EGFP-AP2. Similar to clathrin, the predominant phenotype seen in kymographs is a dramatic increase in long-lived static AP2 structures upon TDD overexpression (Fig. 5A-B, Videos S3-4). AP2 turnover was also impaired in these static structures (Fig. S2D-F). Unlike clathrin, the background intensity of AP2 on the plasma membrane was enhanced by TDD (Fig. 5C-D) and we detected a corresponding increase in association of AP2 with the membrane pellet (Fig. 5E, from 59%±14% for control cells to 76%±14% for cells with TDD overexpression, n=3). These data indicate that TDD can stabilize AP2 complexes on the plasma membrane outside of CCPs.

**Figure 5.**
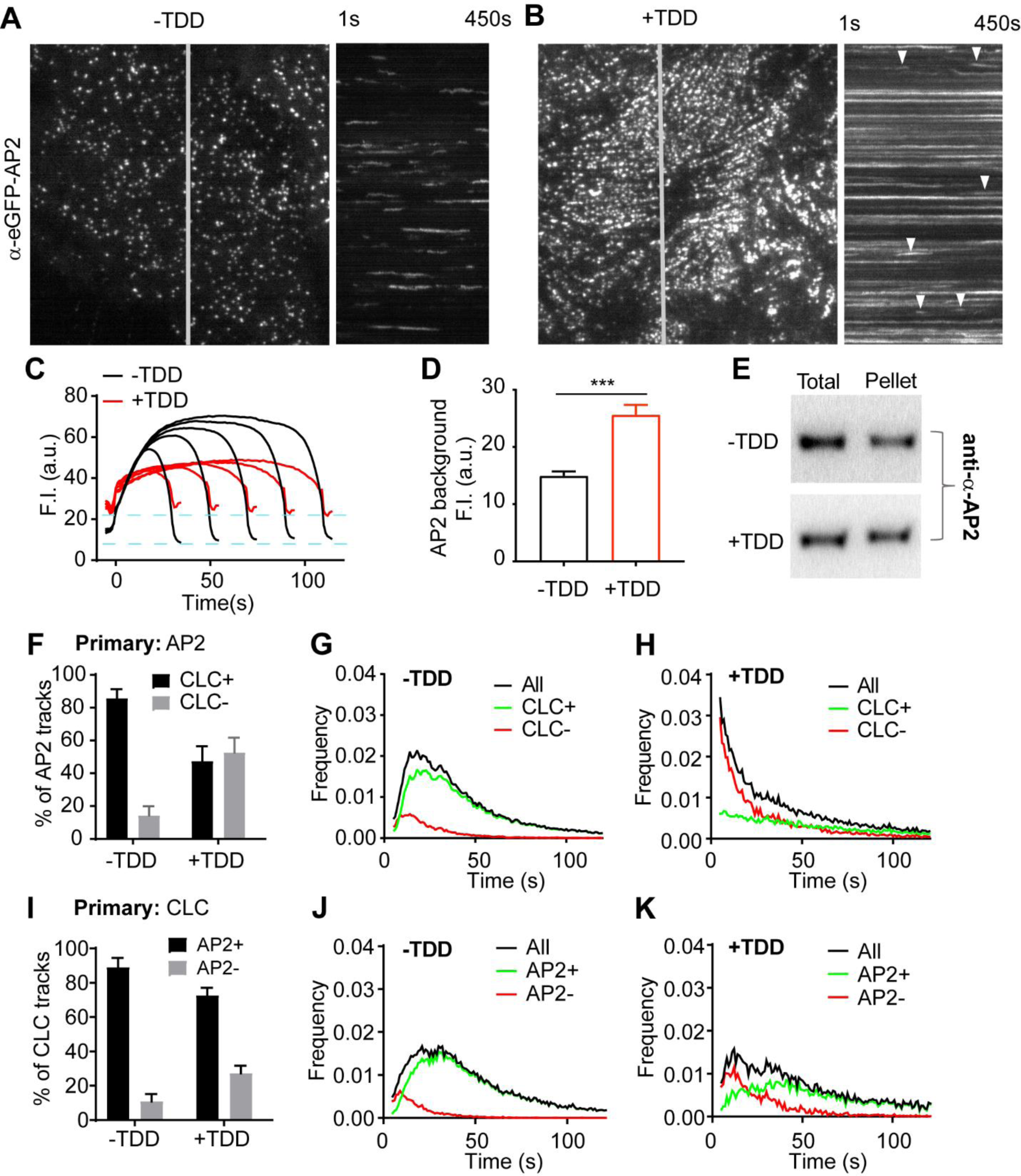
Effects of TDD on AP2-clathrin interactions. (A-D) TIRF imaging of ARPE cells expressing α-eGFP-AP2. (A-B) Single frame from TIRFM movie (7.5min/movie, see Videos S3-4) and corresponding kymographs without (A) and with (B) TDD expression. Arrowheads point to examples of dynamic structures subjected to cmeAnalysis. (C) Mean fluorescence intensity traces of lifetime cohorts of analyzed AP2-labeled structures. Dashed lines indicate the elevation of background fluorescence intensity of AP2 as quantified in (D). Number of analyzed tracks: 105498 for –TDD and 102207 for +TDD. (E) Representative western blot of α-AP2 from subcellular fractionation of ARPE cells with or without TDD expression. (F-K) Dual color TIRFM imaging of ARPE cells expressing both mRuby-CLCa and α-eGFP-AP2. (F-H) Dual color cmeAnalysis was performed using AP2 as primary channel. The percent of AP2 structures with (CLC+) and without (CLC-) detectable clathrin is shown for -TDD and +TDD cells. Number of analyzed tracks: 106310 for control and 77932 for +TDD. Error bars are standard deviations shown in (F). Comparison of lifetime distributions of all detected AP2 structures and those with (CLC+) or without (CLC-) detected clathrin in -TDD (G) and +TDD (H) cells. (I-K) Same as for (F-H), except CLC was assigned as the primary channel. Number of analyzed tracks: 172166 for -TDD and 133604 for +TDD.

Dual-color TIRFM imaging and cmeAnalysis using AP2 (α-eGFP-AP2) as the ‘primary’ detection and clathrin as ‘secondary’ (formerly ‘master/slave’) (Aguet et al., 2013), revealed that clathrin was depleted from a large portion of dynamic AP2 structures upon TDD overexpression (Fig. 5F). AP2 structures that were not stabilized by clathrin were short-lived (Fig. 5G-H). Correspondingly, assigning clathrin as primary, the fraction of AP2-deficient CCPs increased and this subpopulation also was short-lived (Fig. 5I-K). From these data we conclude that TDD inhibits early stages of CME, in part, by disrupting clathrin-AP2 interactions.

### SNX9 depletion phenocopies the inhibitory effects of TDD overexpression

Given that SNX9 interacts with TDD (Fig. 4E), we compared the effects of SNX9 knockdown on CCP dynamics with those of TDD overexpression. As was previously reported (Schöneberg et al., 2017), SNX9 knockdown increased the number of long-lived, static CCPs visible in kymographs (Fig. 6A, Video S5). Analysis of the remaining dynamic structures revealed substantial decreases in the rates of CCP initiation (Fig. 6B) and clathrin recruitment (Fig. 6C-D). Dynamic CCPs were also dimmer (i.e. smaller, Fig. 6E) and, as previously reported (Srinivasan et al., 2018), had prolonged lifetimes (Fig. 6F). Thus, depletion of SNX9 phenocopied many aspects of the effects of TDD overexpression, consistent with overlapping mechanisms of inhibition of CME.

**Figure 6.**
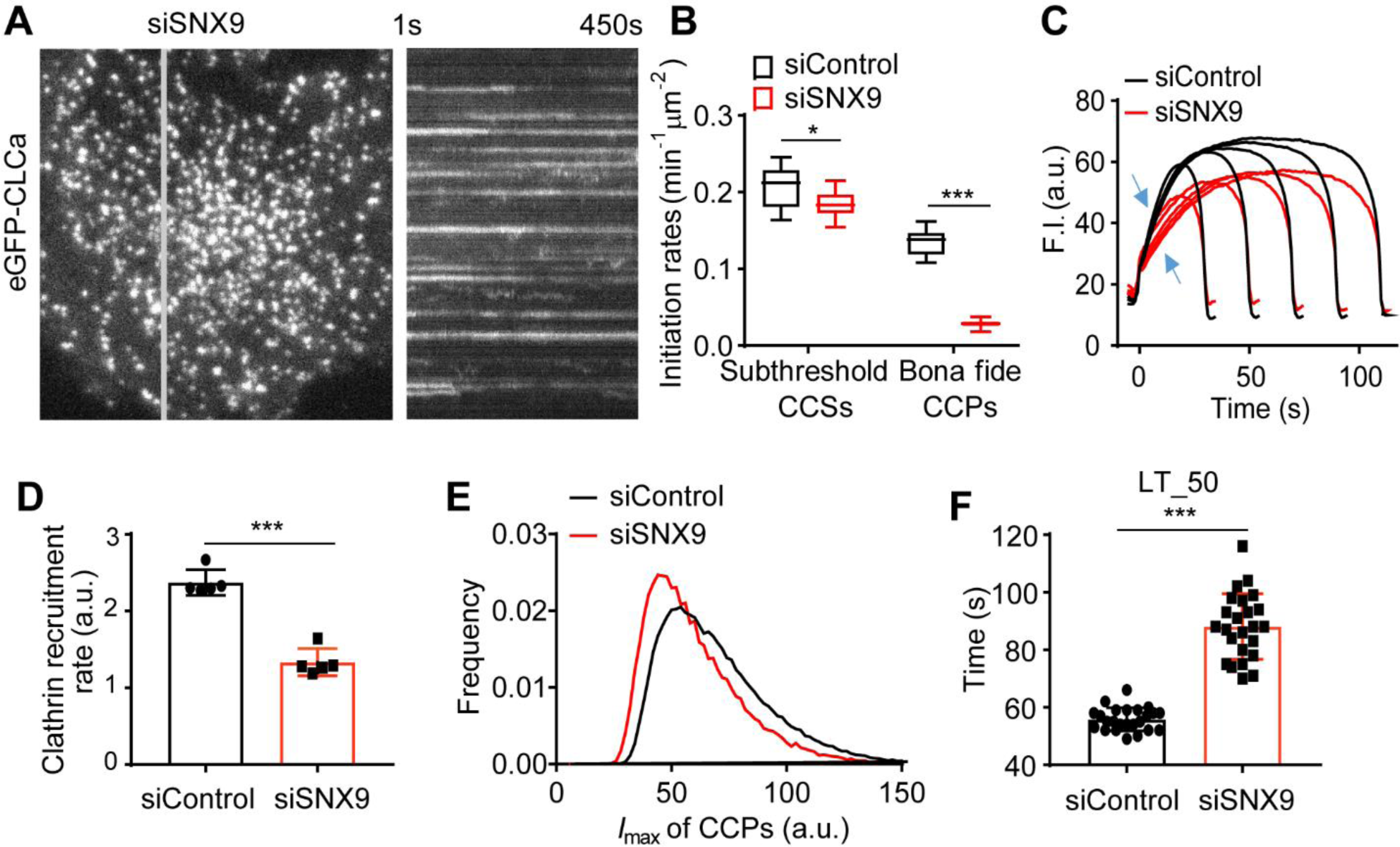
Effects of SNX9 knockdown on CCP dynamics. (A) Single frame from TIRFM movie (7.5min/movie, see Video S5) and corresponding kymograph from region indicated by the gray line of ARPE/HPV cells expressing eGFP-CLCa. (B) Effect of SNX9 knockdown on the initiation rates of *bona fide* CCPs and subthreshold CCSs. (C) Mean clathrin fluorescence intensity traces of lifetime cohorts of analyzed clathrin structures. Growth phase is indicated by blue arrows. (D) Mean slopes of the fluorescence intensity cohorts shown in (C) measured during the growth phase from t = 3s to t = 8s. Each dot represents mean slope of a cohort. (E) Maximum fluorescence intensity (*I*max) distribution of CCPs in control and SNX9 knockdown cells. (F) Medium lifetime (LT_50) of CCPs in control and SNX9 knockdown cells. Number of dynamic tracks analyzed: 294804 for siControl and 152013 for siSNX9. Data ± standard deviations in (B) and (F) are from N=24 movies.

### Design of inhibitory, membrane penetrating peptide-mimetics of TD

TDD overexpression potently and selectively inhibited CME; however, the extended incubation times needed to induce protein expression limited its utility. Moreover, we were interested in dissecting which bindings sites on the TD contributed to its inhibitory effects. Therefore, we designed membrane-penetrating peptides that mimicked the binding sites on TD as potential acute inhibitors for CME, these included the clathrin-box (ter Haar et al., 2000), the W-box (Miele et al., 2004), the a2L site, which also binds clathrin-box motifs (Kang et al., 2009) and the less well-defined site 4 (Willox and Royle, 2012) (Fig. 1B). To this end, six peptides encoding the reported key residues within the four binding sites on TD (Fig. 7A) were designed and synthesized with an N-terminal arginine-rich TAT-sequence for efficient membrane penetration (Copolovici et al., 2014; Walrant et al., 2017) (Fig. 7A). The Clathrin-box1 (designated CBox1), W-box1 (designated Wbox1), a2L, and site 4 peptides were derived from linear 10 amino acid TD sequences (Fig. 7B). Because key residues in the Clathrin-box and W-box binding sites are encoded on discontinuous residues located on adjacent blades (Fig. 7A-B), two additional peptides (Cbox2 and Wbox2) were designed to bring these sequences together in a 10 amino acid peptide engineered to encompass the entire binding motifs.

**Figure 7.**
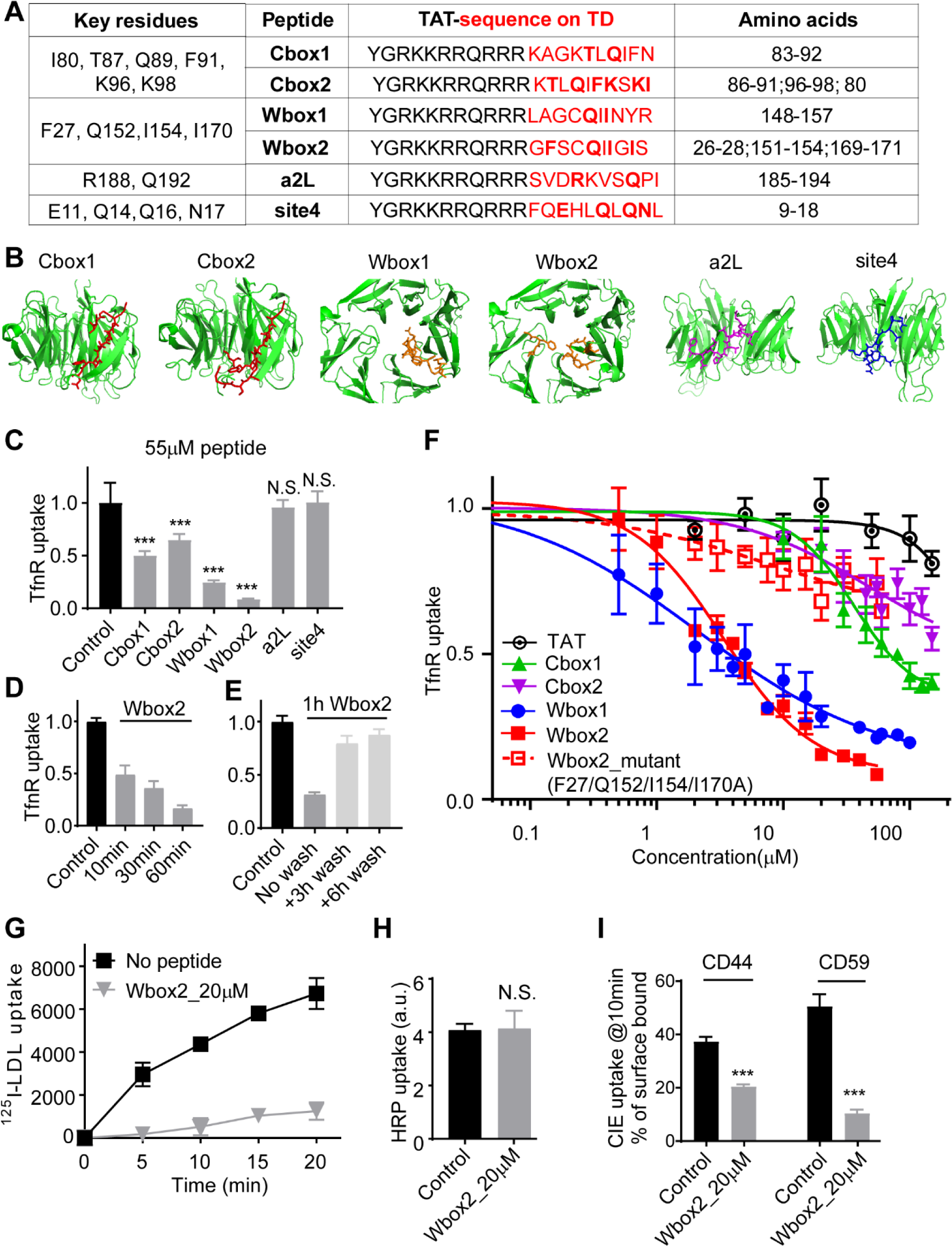
Peptides designed based on TD binding sites inhibit CME. (A) Design of TAT-tagged peptides. Previously reported key residues of the four binding sites in clathrin terminal domain are listed on the left column. Peptides encoded the TAT sequence (black) followed by 10 amino acids derived from TD (red, key residues indicated in bold). Cbox1 (clathrin-box1), Wbox1 (W-box1), a2L and site4 are peptides derived from the linear TD sequences incorporating the key residues. Cbox2 and Wbox2 were designed to incorporate discontinuous key residues located on adjacent blades bringing them together in a 10 amino acid peptide engineered to encompass the entire binding motif. (B) Amino acid sequence of 6 designed peptides are shown as colored sticks on the TD (PDB:1BPO). (C) Comparison of the inhibitory activities of the 6 peptides on TfnR uptake in ARPE/HPV cells. Cells were pre-incubated with peptides (55µM) for 60min at 37°C before uptake assay. (D) Effect of pre-incubation time (10-60min at 37°C, as indicated) on the inhibitory activity of 20µM Wbox2. (E) Reversibility of Wbox2 inhibition. TfnR uptake efficiency was measured in ARPE/HPV cells immediately after a 1h pre-incubation at 37°C with 20µM WBox2 or after removing peptides and incubating cells at 37°C for 3 and 6 additional hours. (F) Relative TfnR uptake efficiency with increasing concentrations of TAT or the indicated TD-derived peptides, as well as Wbox2_mutant after 60min pre-incubation at 37°C. (C-F) The values were normalized to control = 1. Data ± standard deviations are from N=4 replicates. (G) Effect of Wbox2 peptides on ^125^I-LDL uptake in human fibroblasts. Data ± standard deviations are from N=3 replicates. (H-I) Effect of Wbox2 peptides (20μM) on (H) fluid phase uptake of HRP or clathrin-independent endocytosis of (I) CD44 and CD59. Error bars are standard deviations from N=4 replicates.

### Wbox2 is a potent and acute peptide inhibitor of CME

ARPE/HPV cells were pre-incubated for 60 min in the presence of 55µM of each peptide and their effects on TfnR uptake via CME were measured. Both Wbox and Cbox peptides substantially inhibited CME (Fig. 7C); whereas the a2L or site4 peptides did not. Similar to TDD, the inhibitory peptides led to the accumulation of TfnRs on the cell surface (Fig. S4A). Inhibition was rapid and reversible using Wbox2 as an example. More than 50% inhibition was achieved within 10 min of peptide incubation, and CME was largely restored after removing peptides from the culture medium for several hours (Fig. 7D-E). Concentration curves showed more potent inhibition by Wbox1 and Wbox2 (IC50∼3μM), as compared to Cbox1 (IC50∼ 60μM) and Cbox2 (IC50 > 100μM) (Fig. 7F). The TAT peptide, which serves as a negative control, did not inhibit TfnR uptake at concentrations below 100μM. Although their IC50s were comparable, inhibition by Wbox2 exhibited a tighter concentration dependence and achieved a greater degree of inhibition at peptide concentrations >5μM than that of Wbox1. Mutation of the four reported key residues to alanines, greatly reduced the inhibitory activity of the resulting Wbox2_mutant peptide (Fig. 7F), consistent with previous studies showing the importance of these residues for protein interactions (Miele et al., 2004). Importantly, Wbox 2 had no effect on cell viability at concentrations ≤30 µM and exhibited an IC50 (>40 µM) that was 10-fold higher than that needed to inhibit CME (Figure S5).

In addition to the uptake of TfnR and its cargo, Tfn (Fig. S4B-C), Wbox2 also strongly inhibited the uptake of LDL (Fig. 7G), even though it can be supported by the Dab2 CME adaptor independent of AP2(Keyel et al., 2006; Maurer and Cooper, 2006). Moreover, the inhibitory effects of Wbox2 peptides can be extended to other cell types, including normal human fibroblasts, HCC4017 non-small cell lung cancer cells, and HeLa cells (Fig. S4D-F). The Wbox2 peptide did not significantly perturb Tfn recycling (Fig. S4G-H), AP1-mediated export of the mannose-6-phosphate receptor (M6PR) from the Golgi (Fig. S4I-K), or fluid phase uptake (Fig. 7H). However, unlike TDD, Wbox2 significantly inhibited CD44 and CD59 uptake (Fig. 7I). Together these data establish the Wbox2 peptide as a potent, acute and reversible inhibitor of CME.

### Wbox2 inhibits CCP dynamics in a manner similar to TDD

We next investigated the underlying mechanisms of inhibition by Wbox2. Like TDD, application of Wbox2 induced static AP2 and clathrin structures on the plasma membrane (Fig. 8A-B, Video S6), indicative of a late block of CCP maturation. The background intensity of AP2 was also elevated by Wbox2 (Fig. 8C). As was seen with TDD expression (Fig. 5), Wbox2 decreased the fraction of clathrin-positive AP2 structures (Fig. 8D), as well as the fraction of AP2-positive clathrin structures (Fig. 8E) in a concentration-dependent manner. These clathrin-negative AP2 structures and AP2-negative clathrin structures were both short-lived (data not shown). These observations motivated us to test whether Wbox2 inhibits early and late stages of CME by interfering with AP2-clathrin and SNX9-clathrin interactions.

**Figure 8.**
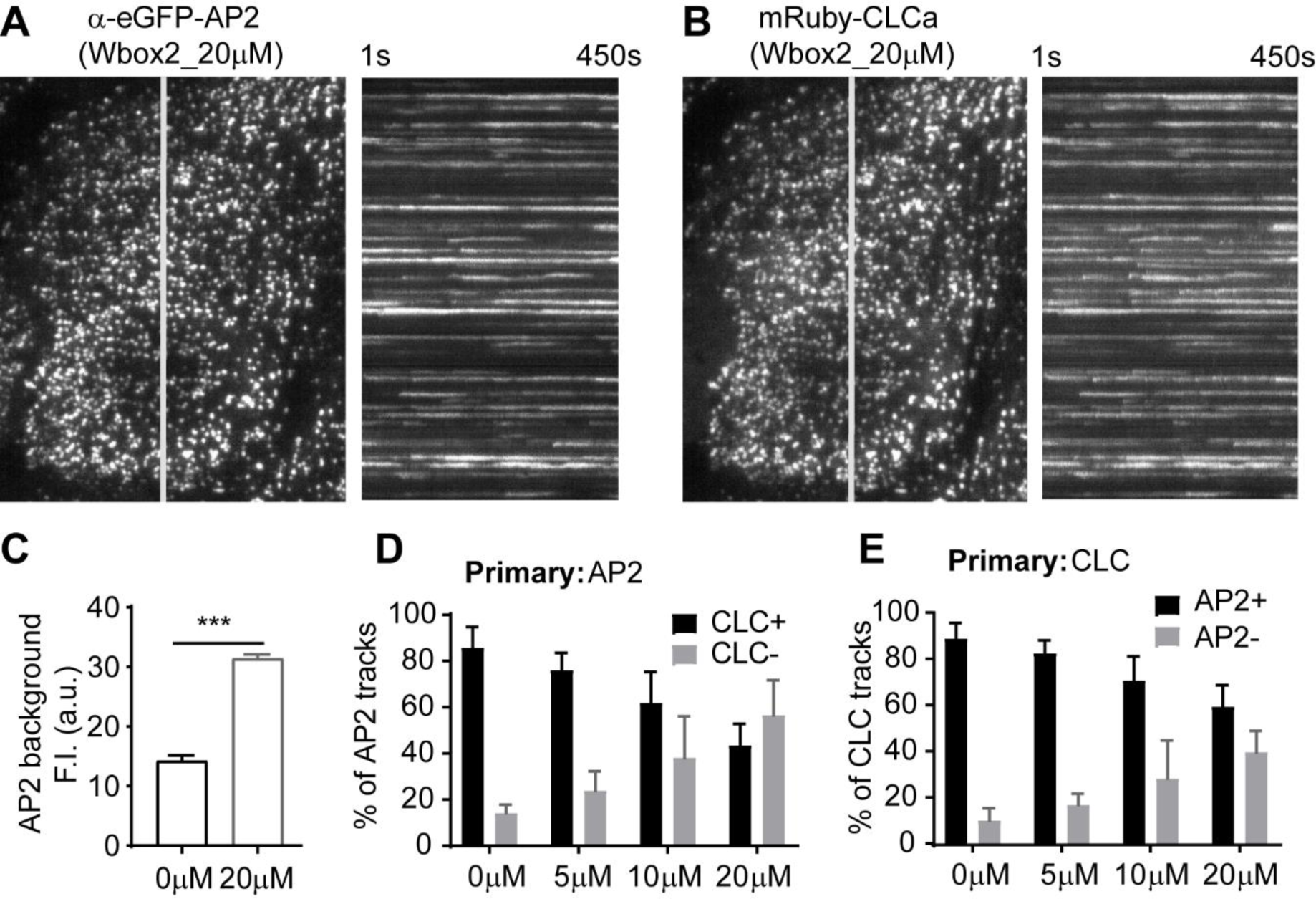
Wbox2 interferes with AP2-clathrin interactions and alters CCP dynamics. Dual color TIRFM imaging of ARPE cells expressing mRuby-CLCa and α-eGFP-AP2 and treated with Wbox2. (A-B) Single frame from TIRFM movie (7.5min/movie, see Video S6) and corresponding kymographs from region indicated by gray line. (C) Background AP2 fluorescence intensity of cells incubated without or with Wbox2 (20μM). (D-E) Dual color cmeAnalysis with either AP2 (D) or CLC (E) assigned as primary channel showing percentage of α-eGFP-AP2 tracks labeled with mRuby-CLCa (D) or mRuby-CLCa tracks labeled with α-eGFP-AP2 (E) determined in the presence of increasing concentrations of Wbox2. Number of dynamic tracks analyzed in (D): 43324 for 0μM, 34664 for 5μM, 33596 for 10μM and 49273 for 20μM. Number of dynamic tracks analyzed in (E): 65192 for 0μM, 42323 for 5μM, 49733 for 10μM and 57758 for 20μM. Error bars are standard deviations.

### Wbox2 perturbs the functions of SNX9 and AP2

Isothermal titration calorimetry (ITC) was used to measure the direct interactions between Wbox2 and purified proteins: SNX9 and hinge+appendage domain of AP2 β subunit (AP2-β-hinge+appendage). Two Wbox2 binding sites were observed for SNX9 with binding affinities of 3.2μM and 44.9μM, respectively (Fig. 9A). Wbox2 was also observed to bind to a fraction of purified AP2-β-hinge+appendage with a binding affinity of 20.1μΜ (Fig. 9A). Given that not all the purified AP2-β-hinge+appendage, showed active binding with Wbox2, potentially due to misfolding or interference by the unstructured hinge, we further investigated whether the addition of Wbox2 inhibits interactions of native AP2 in GST-TD (TD residues: 1-580) pulldown experiments. As shown by others (Schmid et al., 2006), GST-TD efficiently pulled down AP2 from brain lysates. Incubation in the presence of increasing concentrations of Wbox2 reduced the pulldown efficiency of AP2 by GST-TD at concentrations comparable to its effects on CME (Figure 9B-C). Together the ability of Wbox2 to interfere with both clathrin-AP2 and clathrin-SNX9 interactions offers a mechanistic basis for its ability to inhibit both early and late stages of CME.

**Figure 9.**
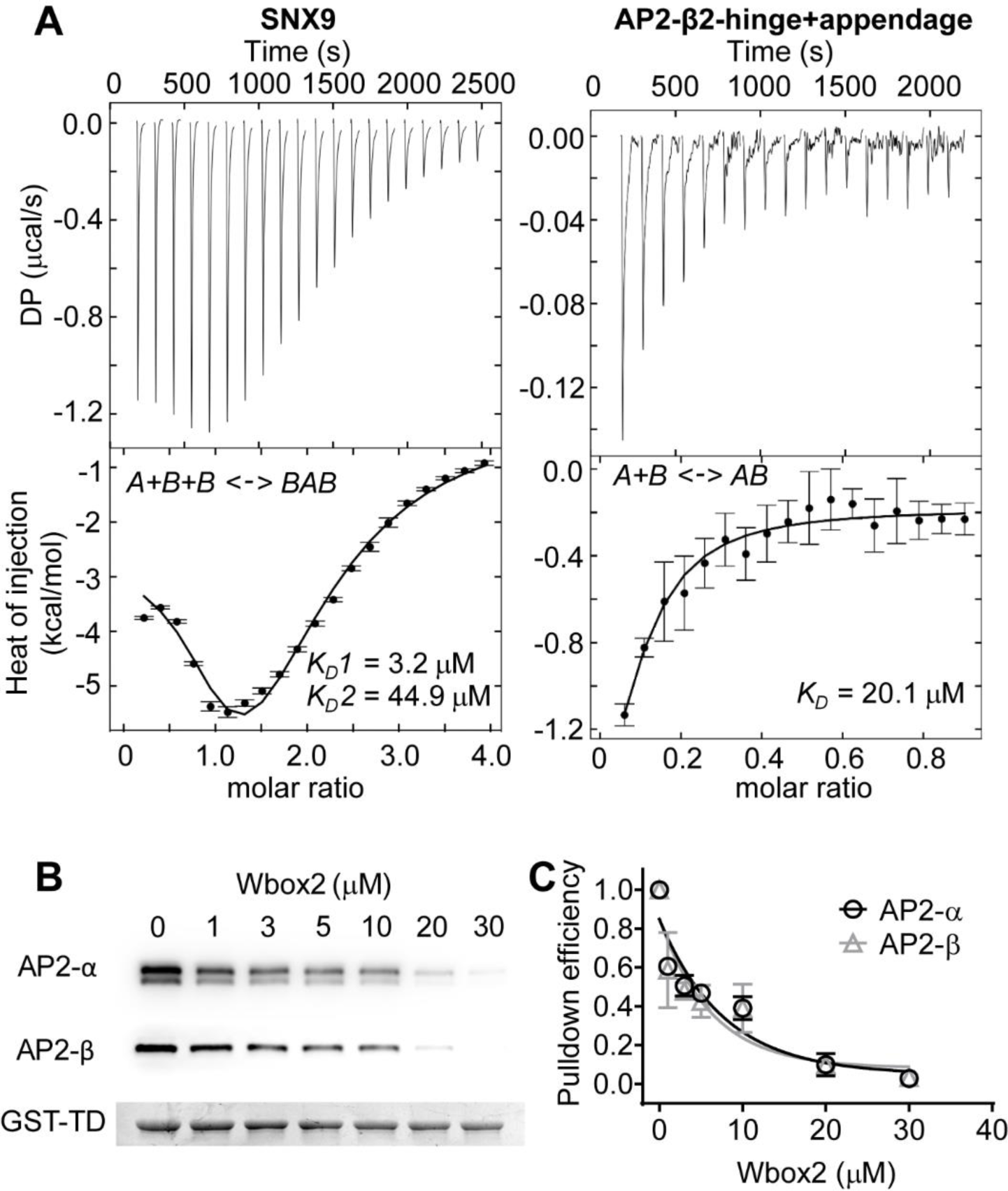
Wbox2 binds to both AP2 and SNX9. (A) ITC measurements of Wbox2 binding to SNX9 and hinge+appendage domain of AP2 β-subunit. Heat curves recorded as a function of time during successive 1.9μl injection of Wbox2 peptide (1.5mM) into the cell containing either SNX9 (80μM) or AP2-β2-hinge+appendage (250μM). Heat curves were executed with NITPIC and fitted with SEDPHAT. The fitting model for SNX9 is A+B+B <-> AB+B <->BAB, with 2 non-symmetric sites, microscop K. The fitting model for AP2-β2-hinge+appendage is A+B <->AB, Hetero-Association. (B-C) GST-TD pulldown of AP2 from mouse brain extract with titrated Wbox2. Representative western blot images are presented in (B) and quantified in (C). GST-TD was used as loading control and each data point represents the average and standard deviations from 3 independent runs.

## Discussion

Coated pit assembly begins when AP2 recruits clathrin to the plasma membrane via interactions between the β-subunit of AP2 and the N-terminal domain (TD) of the clathrin heavy chain (Cocucci et al., 2012; Shih et al., 1995). The seven-bladed β-propeller of TD also binds many EAPs *in vitro* (Dell’Angelica, 2001; Lemmon and Traub, 2012; ter Haar et al., 2000) and is thus seen as a critical hub in the CME interactome (Schmid et al., 2006; Schmid and McMahon, 2007). TD-hub interactions have been proposed to regulate later stages of CME as the coat matures from an AP2-centric hub to a clathrin-centric hub (Schmid et al., 2006; Schmid and McMahon, 2007). However, CME is unaffected by disruption of the major binding sites on the TD, either individually or in combination (Collette et al., 2009; Willox and Royle, 2012). To address this paradox and better define the *in vivo* function of the TD, we characterized the consequences of its overexpression with (TDD) or without (TD) the distal leg, as a competitive inhibitor of TD interactions. TDD/TD overexpression potently inhibited CME and perturbed CCP dynamics, increasing the number of static CCPs and lengthening the lifetimes of the dynamic CCPs that remained. Detailed analysis of the dynamic subpopulation of CCPs revealed an increase in transient, dim CCSs but a reduced rate of initiation of *bona fide* CCPs. This defect in the stabilization of nascent CCPs corresponded to a decrease in the rate and extent of clathrin recruitment. These phenotypes indicate both early (CCP assembly) and late (CCP maturation) roles for TD interactions in CME and provide a mechanistic basis for neurologically diseases linked to frameshift mutations in the gene encoding CHC.

TDD overexpression potently inhibited CME without affecting bulk fluid-phase endocytosis or the clathrin-independent uptake of CD44 or the GPI-anchored protein CD59. Thus, these studies establish TDD as a new dominant-negative construct able to specifically inhibit CME.

Co-IP identified the AP2 complex and SNX9 as major binding partners of TDD, which together can account for both the early and late effects of TDD overexpression. TDD overexpression reduced the extent of colocalization of AP2 with clathrin and stabilized AP2 on the PM outside of CCPs. Both clathrin-deficient AP2 clusters and AP2-deficient clathrin clusters were short-lived, indicating a role for TD-AP2 interactions in stabilizing nascent CCPs. Similarly, SNX9 knockdown phenocopied many of the effects of TDD overexpression, including the formation of static clathrin structures, decreased stabilization of CCPs, slower rates of clathrin recruitment, and prolonged lifetimes of dynamic CCPs. Thus, the inhibitory effects of TDD on CME can be accounted for by its interference with clathrin-AP2 and clathrin-SNX9 interactions (Fig 1C, Mechanisms 2 and 3). It was surprising that TDD was not incorporated into CCPs, given that TDD encodes both AP2 binding and clathrin assembly domains. These findings are consistent with the hypothesis that a high degree of cooperativity involving simultaneous interactions with two AP2 complexes is required to initiate clathrin recruitment (Cocucci et al., 2012; Moskowitz et al., 2005).

Our inability to detect interactions between TDD and most EAPs reported to bind TD *in vitro*, including amphiphysin, eps15, AP180 or epsin1 (reviewed in (Lemmon and Traub, 2012; Royle, 2006; Schmid and McMahon, 2007)), is consistent with previous pull-down assays from bovine brain extracts using purified GST-TD (Schmid et al., 2006). It is possible, as previously suggested (Lemmon and Traub, 2012; Royle, 2006; Schmid and McMahon, 2007), that EAPs bind TDD with low affinity and only associate with CCPs when clathrin assembles into a lattice creating opportunities for high avidity interaction.

Unexpectedly, AP1 did not robustly co-precipitate with TDD (Table S2), despite an overall sequence identity of 83% and conservation in the clathrin-box binding motifs of the β-subunits of AP1, AP2 and AP3 (Ahle and Ungewickell, 1989; Gallusser and Kirchhausen, 1993; ter Haar et al., 2000). Consistent with this, trafficking from the Golgi, which relies upon AP1-clathrin interactions, was not significantly perturbed by TDD overexpression. These findings bring into question the conserved role of clathrin-box sequences alone in mediating AP-clathrin interactions. Indeed, mutation of the clathrin-box in the β-subunit of AP3 does not impair AP3 function (Peden et al., 2002). Similarly, a clustered β3 hinge-ear construct is unable to recruit clathrin to PM, in contrast to an identical β2-hinge-ear construct that was used to ‘hot-wire’ CME (Wood et al., 2017). The stronger affinity of AP2 for clathrin may reflect the bipartite nature of AP2-clathrin interactions due to the existence of a second clathrin binding site on the β2-appendage domain (Edeling et al., 2006; Owen et al., 2000). Our findings suggest a need for further exploration of the interactions of AP2, as well as other AP complexes with clathrin.

To overcome the limitations of long-term overexpression of TDD and to probe the contributions of different TD binding sites towards its inhibitory activity, we designed peptide-based mimetics of the individual binding surfaces on the TD. Both Wbox- and Cbox-mimicking peptides significantly inhibited CME. Of these, the Wbox peptides were most potent suggesting, unexpectedly, that the W-box site interactions play a more important role in CME, although we cannot rule out the possibility that our Cbox peptide designs do not optimally mimic the proper folding of this interaction surface. Further peptide design, including for a2L and site 4 mimics, may be warranted.

Strikingly, treatment of cells with Wbox2, the most potent Wbox-derived peptide, reproduced many of the phenotypes resulting from overexpression of TDD, providing further evidence for the importance of the W-box site in CME. This region of TD is reported to bind proteins via a PWxxW motif, which has only been identified in amphiphysin and SNX9 (Lemmon and Traub, 2012; Miele et al., 2004). We did not detect amphiphysin in any of our pulldown experiments (Table S2); thus, we ascribe some of the effects of Wbox2 on competition of TD-SNX9 interactions, which plays multiple roles in CME (Schöneberg et al., 2017; Srinivasan et al., 2018). Indeed, Wbox2 binds directly to two sites on SNX9 with affinities comparable to its inhibitory effects on CME. Given that the clathrin-box site is the presumed binding site for AP2, it was surprising that Wbox2 also phenocopied TD in its ability to interfere with AP2-clathrin interactions. This was confirmed *in vitro* as we could detect direct binding of Wbox2 to the β2-hinge-ear and found that it inhibited pulldown of AP2 complexes by GST-TD, again with affinities comparable to its inhibitory effects on CME.

Given the β-propeller fold of the TD, the strong inhibitory activity of Wbox2 was somewhat surprising. Several factors can explain these findings. First, the key residues demarking the W-box site are located on blades 1, 5 and 6 of the TD β-propeller (Miele et al., 2004). Thus the β-propeller fold serves to generate a central interaction surface, which is presumably mimicked by our Wbox2 design. Second, the W-box site spans the top and center of the TD (Fig. 1B), and is thus positioned to take greatest advantage of the bipartate clathrin-interaction surfaces on the β2-hinge and ear (Lemmon and Traub, 2012; Willox and Royle, 2012). Finally, the W-box site faces the membrane in an assembled coat (Fotin et al., 2004), and may thus be better positioned to interact with EAPs involved in cargo recognition or membrane remodeling.

Clathrin heavy chain variants bearing mutations that disrupt either the clathrin-box, the W-box or both appear to be fully functional in cells (Collette et al., 2009; Willox and Royle, 2012). These observations suggest that AP2 and EAPs can interact with functionally redundant sites on clathrin. Clathrin-binding motifs are promiscuous (Lemmon and Traub, 2012) and often occur in multiple copies within intrinsically disordered regions (IDRs). The clathrin-binding motifs of IDRs have been proposed to bind to clathrin lattices via a “line-fishing” mechanism involving dynamic and non-specific association/dissociations without protein folding (Zhuo et al., 2010). The Wbox and Cbox peptides could bind to and mask multiple sites on TD-binding partners and thus more globally disrupt these interactions.

That Wbox2 interferes with both AP2- and SNX9-clathrin interactions provides a mechanistic basis for its ability to phenocopy overexpression of TDD and inhibit both early and late stages of CME. However, unlike TDD, Wbox2 inhibited CD44 and CD59 endocytosis. There are two possibilities for this unexpected difference. First, it seems likely that cargo such as CD44 and CD59, which have no known interactions with cytoplasmic adaptor proteins would freely diffuse on the cell surface and be captured by endocytic coated pits. Indeed, Nichols and colleagues showed that in unperturbed HeLa, COS and RPE cells >90% of CD59 was taken up in nascent clathrin coated vesicles together with Tfn (Bitsikas et al., 2014). That their uptake is inhibited under acute, but not prolonged conditions, could be explained by previous studies showing that clathrin-independent endocytic pathways are upregulated following prolonged inhibition of CME, for example by expressing dominant-negative proteins or siRNA knockdown of clathrin or AP2 (Bitsikas et al., 2014; Damke et al., 1995). Consistent with this, CD44 uptake is not inhibited by dominant-negative dynamin overexpression (Mayor et al., 2014); but has been shown to be inhibited following acute treatment with dynasore (Takahashi et al., 2015). Similarly, acute treatment with Pitstop, inhibits both CME and CIE (Dutta et al., 2012). The second possibility relates to the ability of Wbox2 to bind two sites on SNX9 with high affinity. Previous studies have shown that siRNA knockdown of SNX9 inhibits CD44 uptake (Bendris et al., 2016) and that SNX9 colocalizes with PM-derived tubules bearing GPI-cargo (Yarar et al., 2007). Thus, Wbox2 could be inhibiting these cargo-selective pathways of CIE, while not pertubing bulk fluid phase uptake. However because TDD also binds SNX9 we favor the former hypothesis, athough further studies are necessary and caution should be taken in interpreting results when using these reagents.

While, further studies are underway to design new, more selective peptide-based inhibitors, Wbox2 provides a significant advance over existing chemical inhibitors of CME. Specifically, both dynasore and Dyngo-4a have been shown to have numerous off-target effects unrelated to CME (Park et al., 2013; Persaud et al., 2018) and Pitstop is cytotoxic at concentrations only ∼2-fold higher than its reported inhibitory effects on CME (Rosselli-Murai et al., 2018). In contrast, our structure based design, together with biochemical characterization of Wbox2 binding and inhibition and control experiments establishing that a peptide bearing mutations known to perturb the W-box interface on TD is not any longer inhibitory, provide a strong basis for the on-target effects of this peptide.

In summary, we show that TDD over-expression inhibited CME primarily through interference of clathrin interactions with SNX9 and the AP2 complex. These interactions were required at multiple stages during CME. We further generated TAT-tagged membrane-penetrating peptides based on binding sites on the TD and identified a peptide mimic of the W-box site, Wbox2, that potently and acutely inhibited CME, phenocopying the effects of TDD overexpression. Importantly, we detailed the mechanism of Wbox2 inhibition, which accounts for its early and late effects on CCP maturation.

## Materials and Methods

### Generation of constructs and viruses

#### TDD constructs

The TDD fragment (residues 1-1074) in a pET23d vector was kindly gifted by Dr. Frances Brodsky (UCL, London, UK), and cloned by seamless technique into a pADT3T7tet vector with an HA tag at the N-terminus. TD was generated by mutating T495 to a stop codon using the TDD construct as template.

#### Adenovirus generation

Recombinant adenoviruses were generated as previously described(Damke et al., 2001; Kadlecova et al., 2017). Briefly, cDNA containing target protein sequence TDD, TD or tTA was transfected into CRE4 HEK293 cells together with ψ5 DNA. Viruses were harvested by 2 cycles of freeze-thaw and stored at -80°C. The harvested viruses were used to infect more HEK293 cells for virus expansion.

#### Peptide design, synthesis and application

Peptides encoding the TAT sequence (YGRKKRRQRRR) followed by 10 amino acid sequences covering the four reported binding sites on the clathrin terminal domain (see Fig. 7A) were synthesized by GenScript (Piscataway, NJ) with > 95% purity. Peptides were dissolved in dH2O and further diluted in PBS4+ (1X PBS buffer with addition of 0.2% bovine serum albumin, 1mM CaCl2, 1mM MgCl2, and 5mM D-glucose) or cell culture medium (for imaging) when applied to cells. Peptide solutions were stored at -20°C.

### Cell culture, viral infection and siRNA knockdown

ARPE19 (herein called ARPE) and ARPE19/HPV16 (herein called APRE/HPV) cells were obtained from ATCC and cultured in DMEM/F12 medium with 10% FBS. Expression of fluorescent protein-labeling CLCa and/or α subunit of AP2 was achieved by infection with lentiviruses carrying a pLVX-puro vector (mRuby2-CLCa), a pMIB6 vector (eGFP-CLCa) or a modified pMIB6 vector lacking IRIS and BFP expression (α-eGFP-AP2, eGFP is encoded within the linker region of alpha subunit). Stable cell lines expressing fluorescent tags were sorted by FACS after 72 hours. HeLa cells were cultured in DMEM medium with 10% FBS. Human Fibroblast cells were cultured in low glucose DMEM medium with 10% FBS. HCC4017 cells were cultured in RMPI medium with 5% FBS. All cell lines were cultured at 37°C in 5% CO2.

TDD, TD and tTA adenoviruses were applied to cells directly in culture medium. Pilot experiments were conducted to optimize the volumes of each virus preparation need for uniform infection (see Fig. S1A-B). Experiments were conducted after 12-18 hours, allowing for adequate viral infection and protein expression. Protein expression could be regulated by adjusting tetracycline concentration. All experiments were performed in the absence of tetracycline allowing for maximum protein expression except for Fig. 1A-C.

For knockdown of SNX9, transfection of siRNA (Sigma, pool of two siRNAs: UAAGCACUUUGACUGGUUAUU and AACAGUCGUGCUAGUUCCUCA) was conducted in Opti-MEM with Lipofectamine RNAi-MAX (Life Technologies). Cells were plated on a 6-well plate (200,000 cells/well) and transfected with siRNA after attaching to the plate. For transfection, 100μl Opti-MEM with 6.5µl Lipofectamine RNAi-MAX and 100μl Opti-MEM with 110pmol siRNA were incubated at room temperature for 5min, the two reagents were then mixed and further incubated at room temperature for 10min. The mixture was then added dropwise to cells. Two rounds of transfection were carried out through 5 days to achieve over 90% target protein knockdown.

### Endocytosis (uptake) assay

Internalization of TfnR, CD44, CD59 and Biotin-Tfn was quantified by in-cell ELISA following established protocols(Conner and Schmid, 2003; Srinivasan et al., 2018). Cells were seeded in 96-well plates (Costar) at 60-70% confluency and grown overnight. Before the assay, cells were starved for 30min in PBS4+ at 37°C or treated with peptides in PBS4+ at 37°C for the indicated time points. After starvation, cells were moved to 4°C and culture media was replaced with cold PBS4+ containing: 5μg/ml HTR-D65 (anti-TfnR mAb)(Schmid and Smythe, 1991), anti-human CD44 or anti-human CD59 (BD Pharmingen) or 5μg/ml Biotin-Tfn (Sigma). For single round assay, the above reagents were kept with cells for 20min at 4°C and then washed out prior to uptake assay. For multi-round assays, the above reagents were kept with cells throughout the uptake process. In parallel, some cells were kept at 4°C for the measurement of surface-bound ligands and blank controls while some were incubated in 37°C water bath for the indicated time points. Acid wash (0.2M acetic acid, 0.2M NaCl, pH 2.3) was used to remove surface-bound antibodies, followed by washing with cold PBS. All were fixed with 4 % paraformaldehyde (PFA) (Electron Microscopy Sciences, PA) in PBS for 30min at 37°C. 0.1% Triton X-100 was added for 6.5min to permeabilize cells followed by addition of blocking buffer. HTR-D65, CD44 and CD59-treated cells were blocked with Q-PBS (PBS, 2% BSA, 0.1% lysine, 0.01% saponin, pH 7.4) while Biotin-Tfn-treated cells were blocked with 2 % casein. Surface-bound and internalized HTR-D65, CD44 and CD59 were probed with goat-anti-mouse antibody conjugated with HRP (Sigma-Aldrich), while Biotin-Tfn was probed with Streptavidin-POD conjugate (Sigma). Color was developed using OPD solution (Sigma-Aldrich) and absorbance was read at 490 nm (Biotek Synergy H1 Hybrid Reader). Cell number variation among wells was accounted for by BCA assay reading at 562 nm.

Fluid phase uptake assays were conducted using horseradish peroxidase (HRP, Sigma) as the readout. Cells were seeded in 96-well plates (Costar) at 60-70% confluency and grown overnight. Before assay, cell culture medium was changed to PBS4+ and cooled down to 4°C. Ice-cold HRP solution (1mg/ml) was then added to cells and moved to 37 °C water bath for 30 min. In parallel, some cells were kept at 4°C as blank control. All the cells were acid washed (0.2M acetic acid, 0.2M NaCl, pH 2.3) and lysed with ELISA blocking buffer (1% TX-100, 0.1% SDS, 0.2% BSA, 50mM NaCl, 1 mM Tris, pH 7.4) for 1h at 4°C. Subsequently, Color was developed using OPD solution (Sigma-Aldrich) and absorbance was read at 490nm (Biotek Synergy H1 Hybrid Reader).

^125^I-LDL uptake was conducted as previously described(Lombardi et al., 1993; Michaely et al., 2007). Briefly, cells were treated or not with peptide at the indicated concentrations for 30min in Medium A (D-MEM supplemented with 20mM HEPES pH=7.5 and 10% lipoprotein poor serum) in a 37°C/5%CO2 incubator. Cells were then washed with ice cold Medium B (Bicarbonate free MEM supplemented with 20mM HEPES pH=7.5 and 10% lipoprotein poor serum) and incubated with 20μg/ml ^125^I-LDL with or without the indicated concentration of peptide in Medium B for 1hr at 4°C. Medium was replaced with warm (37°C) Medium and cells were incubated for the indicated times after which cells were extensively washed with ice cold PBS and incubated with 1 mg/ml Protease K in Buffer A (1xPBS, 1mM EDTA) for 1hr at 4°C. The cell suspension was then centrifuged at 5000xg for 10min over a cushion of 10% sucrose in PBS. The tubes were frozen in liquid nitrogen, cut to separate the cell pellet (internal) from the solution (surface bound material released by protease K) and counted on a gamma counter.

### Recycling Assay

Biotin-Tfn recycling was conducted following previous protocols (Chen et al., 2017). Cells, which were cultured in biotin-free media for 48hrs, were seeded in 96-well plates (Costar) at 60-70% confluency and grown overnight. Before assay, cells were starved for 45 min in PBS4+ at 37°C or treated with peptides in PBS4+ at 37°C for indicated time points. After starvation, cells supplied with 100μl 5μg/ml Biotin-Tfn in PBS4+ for 10 min or 30 min to load Biotin-Tfn into cells. When loading was complete, cells were cooled down to 4°C and washed with PBS4+ to remove extra Biotin-Tfn. Some cells were kept as ‘loading’ control in 4°C. The remaining cells were first treated with avidin and biocytin to mask the surface-bound Biotin-Tfn, and then incubated at 37°C for the indicated times. All the cells were then acid washed and fixed with 4%PFA as before. Blocking steps were the same as the Biotin-Tfn uptake assay. Data were expressed as % of total intracellular ligand remaining relative to the initial load.

### HA-TDD immunoprecipitation

ARPE cells in a 15cm dish of about ∼ 90% confluency were infected with adenoviruses and incubated for >15hrs at 37°C in 5% CO2 to allow for HA-TDD overexpression. Then cells were detached with 5ml 50mM EDTA at 37°C for 10 min and neutralized with 0.2ml 1M MgCl2. After spinning down cells at 500xg, 4°C for 3min, cells were washed with 1ml ice-cold PBS to remove residual EDTA and MgCl2. Cells were then spun down again and resuspended in 1ml HEPES/KCl buffer (25mM HEPES, 150mM KCl, 1mM MgCl2, 1mM EGTA, 1x protease inhibitor + 1x phosphatase inhibitor, pH=7.4) + 0.5% Triton X-100. With 30min rotation in cold room and occasional vortex, the cells were lysed and spun at 500xg, 4°C for 3 min to remove nuclei. Subsequently, 10μl HA antibody (0.5-0.7mg/mL, Sigma) was added to cell lysate containing 0.5g proteins (determined by Bradford assay) and 0.25% Triton-X100 (adjusted by adding equal volume of HEPES/KCl buffer without Triton X-100). The reaction was allowed to proceed by rotation at 4°C for 2hrs and then immunoprecipitated with 60μl Protein G Sepharose (Sigma). The sediments were washed twice with HEPES/KCl buffer + 0.25% Triton-X100. The final samples were heat denatured and run into SDS-page gel before sending to the proteomics core facility for further sample preparation and MassSpec analysis. The identified protein IDs and abundances were output as a spreadsheet for each sample and Reactome Pathway Database was used for pathway analysis (See Supplemental Tables S1 and S2).

### Subcellular fractionation

Confluent ARPE or ARPE/HPV cells in a 10cm dish were detached with 3ml 50mM EDTA at 37°C for 10min. Extra EDTA after detachment was neutralized with 0.2 ml of 1 M MgCl2. Cells were then spun down at 500xg for 3min and washed with cold PBS. After another spin to remove PBS, cells were resuspended in 0.5ml ice-cold lysis buffer (25mM HEPES, 250mM sucrose, 1mM MgCl2, 2mM EGTA, pH=7.4). Cells were then lysed by 5 rounds of freeze-thaw cycle (rapid freezing in liquid nitrogen and slow thawing in room temperature water bath, followed by brief vortex and 10s sonication). Intact cells and nuclei were removed by centrifugation for 2min at 500xg. 0.1ml lysed sample was kept as loading total, and the rest sample (0.4ml) underwent ultracentrifugation at 4°C for 30min at 110,000xg to separate membranes (pellet) from cytosol (supernatant). Supernatant was transferred to another low binding eppendorf, and the pellet was resuspended with the same volume of lysis buffer (0.4ml). Subsequently, total, pellet and supernatant were heat denatured in 1X sample buffer and probed by western blotting.

### TCV purification

Assembled clathrin/AP2 coats (TCVs) were isolated following cell lysis in 0.5% TX-100, as previously described(Pearse, 1982). Briefly, confluent ARPE/HPV eGFP-CLCa cells in five 15cm dishes were detached with 50mM EDTA (5ml each dish) at 37 °C for 5-10min. Subsequently, extra EDTA was neutralized with 1ml 1M MgCl2. Cells were then collected at 500xg for 3 min and washed with cold PBS. PBS removal was followed by resuspension in 2 ml ice-cold lysis buffer (100mM Mes, 0.2mM EGTA, 0.5mM MgCl2, 0.5% Triton X-100, pH 6.2). The resuspended cell lysates were split into 2 low binding Eppendorf tubes and rotated at 4°C for 30min with occasional vortex. Lysed cells were spun at 500xg, 4°C for 3min to remove nuclei before ultracentrifugation at 121,000xg, 4°C for 45 min to collect enriched TCVs in the pellet.

### Immunofluorescence

Cells seeded on gelatin-coated 22x22mm glass (Corning, #2850-22) were rinsed 3x2ml with PBS and then fixed with 4% PFA (Electron Microscopy Sciences, PA for 30min at 37°C. Subsequently, 0.5% TX-100 was applied to permeabilize the cells. After blocking with Q-PBS, primary antibody and secondary antibodies were added consecutively to cells and incubated for 1hr at room temperature with thorough washes in between. Cells were mounted up on cover slides with spacers and sealed with PBS for TIRF or wide filed imaging.

### TIRFM live cell imaging and CCP quantification of dynamics

Cells were seeded on a gelatin-coated 22x22mm cover glass (Corning, #2850-22) overnight and moved to fresh cell culture medium 30mins before being mounted to a 25x75mm slide (Thermo Scientific, #3050). Imaging was conducted with a 60×, 1.49-NA Apo TIRF objective (Nikon) mounted on a Ti-Eclipse inverted microscope. Perfect focus was used during time-lapse imaging. TIRF penetration depth was∼80nm. Videos were acquired for 7.5min at the rate of 1 frame/s for all live cell imaging. For dual color-imaging, two sequential images of primary and secondary channel were taken within 1s and the video length is the same as single channel imaging. Published cmeAnalysis software was used for CCP detection, tracking and quantification(Aguet et al., 2013; Jaqaman et al., 2008; Loerke et al., 2011). More than 10 cells were imaged per condition and the number of total analyzed tracks are indicated in Figure captions.

### TEM imaging

Cells grown on gelatin-coated glass-bottomed MatTek dishes were washed with cold PBS and fixed in 2.5% (v/v) glutaraldehyde in 0.1M sodium cacodylate buffer. Processing for embedding and sectioning continued as follows: after three rinses in 0.1M sodium cacodylate buffer, they were post-fixed in 1% osmium in 0.1M sodium cacodylate buffer for 1h. Cells were rinsed in 0.1M sodium cacodylate buffer and *en bloc* stained with 0.5% tannic acid in 0.05M sodium cacodylate buffer for 30min. After two rinses in 1% sodium sulfate in 0.1M sodium cacodylate buffer, samples were rinsed three times in 0.1M sodium cacodylate buffer and five times in water. Samples were then dehydrated through a series of increasing concentrations of ethanol, then infiltrated and embedded in Embed-812 resin. Enough resin was added into the MatTek dishes to just fill the well and polymerized at 60°C. Polymerized samples were dropped into liquid nitrogen to pop out the resin disks from center of the MatTek dish. Two resin disks containing the same sample were sandwiched together with fresh Embed-812, monolayers were facing each other. Resin disks were polymerized at 60°C overnight and sectioned along the longitudinal axis of the two monolayers of cells with a diamond knife (Diatome) on a Leica Ultracut UCT 6 ultramicrotome (Leica Microsystems). Sections were post-stained with 2% (w/v) uranyl acetate in water and lead citrate. Imaging was done on a JEM-1400 Plus transmission electron microscope equipped with a LaB6 source operated at 120kV using an AMT-BioSprint 16M CCD camera. Purified TCVs were applied to 200-mesh copper grids and allowed to adsorb for 10min, after which the grids were dipped in water droplets to wash away un-adsorbed material. The grids were then incubated in 2% uranyl acetate for 2min. After 3 dips in water droplets, grids were dried and imaged.

### Protein purification and Isothermal titration calorimetry (ITC) measurements

GST-SNX9 in a pGEX-KG vector and His6x-β2-hinge+appendage in a pET vector (gifted from Dr. Linton Traub, Univ. of Pittsburgh) were expressed in BL21(DE3) and then affinity-purified using glutathione Sepharose 4B beads (ABT) and HisTrap HP column (GE Healthcare), respectively. The affinity-purified GST-SNX9 was thrombin-digested to remove GST. The resulting SNX9 and His6x-β2-hinge+appendage proteins were separately applied to a HiLoad 26/600 Superdex 200 pg column (GE Healthcare). Peak fractions of target proteins were collected and concentrated using Amicon Ultra-15 10K Centrifugal filters (Sigma-Aldrich).

Binding of Wbox2 to SNX9 and AP2-β2-hinge+appendage domain was investigated by ITC using a MicroCal iTC200 (GE Healthcare Life Sciences). Measurements were performed in 20mM HEPES, 150mM NaCl, 1mM TCEP, pH=7.4 and at 20°C. Protein and peptide concentrations were determined by absorbance at 280nm and 205nm, respectively. The peptides were injected from a syringe in 21 steps up to a molar excess over the protein concentration. Titration curves were executed with NITPIC and fitted with SEDPHAT. GUSSI was further used to output the figures presented.

### GST-TD pulldown from mouse brain extract

A mouse brain was homogenized using a dounce tissue grinder (Sartorius) in 1ml MES buffer (100mM MES, 1mM EGTA, 0.5mM MgCl2, 0.1mM PMSF, pH=6.4). After spinning at 5000xg, 4°C for 3min, the supernatant was ultracentrifuged at 110,000xg for 60min. The resulting pellets was then resuspended in extraction buffer (3 vol 1M Tris pH 8 : 1 vol Mes buffer, add PMSF and DTT to 0.1mM each) for 30min at room temperature. The resuspended solution was ultracentrifuged at 110,000xg for 60min again to collect supernatant containing released coat proteins. The brain extract buffer was exchanged to pulldown reaction buffer (50mM Tris, 150mM NaCl, 0.1mM EGTA, pH=6.4) using zeba spin desalting columns (ThermoFisher Scientific).

In pulldown assay, 75μg GST-TD (residues 1-580) in reaction buffer was mixed with coat protein fractions (10% of total extract from a mouse brain) and indicated concentration of Wbox2 peptide. The mixture was incubated with 50μl 50% slurry of glutathione Sepharose 4B beads (ABT) and left rotating overnight at 4°C. Subsequently, beads were collected by spinning down at 1000xg for 2min and washed twice with reaction buffer. The beads were resuspended in Laemmli sample buffer and denatured at 95°C before running on 7.5% SDS-PAGE and western blotting.

### Quantification and statistical analysis

Endocytosis and recycling assays were performed in biological replicates. TIRFM data was biologically reproduced and a representative data set from the same day was presented. The intensity of protein blots was analyzed with ImageJ. For all the data, error bars are standard deviations and the statistical significance was analyzed by two-tailed Student’s t-test. ∗, p≤0.05; ∗∗, p≤0.01; ∗∗∗, p≤0.001.

## Data availability

Mass Spec data that support the findings of this study is available in Supplementary Tables S1 and S2. All other data supporting the findings of this study are available from the corresponding author on reasonable request.

## Code availability

The software and algorithms for data analyses used in this study are all well-established from previous work and are referenced throughout the manuscript. No custom code was used in this study.

## Reagents

A list of reagents used in this study is supplied as Supplementary Table S3.

## Supplemental material

The supplemental materials include 5 supplemental figures and their corresponding legends, 3 supplemental spreadsheet tables containing the information of proteins identified by co-IP and MassSpec, 6 supplemental videos and their corresponding legends.

## Acknowledgements

We thank Dr. Frances Brodsky for reagents. We thank the UTSW proteomics core facility for help with sample processing and analysis, the Molecular Biophysics Core facility for the help with ITC and the Electron Microscopy Core facility for expert assistance in processing EM samples. This work is supported by the Welch grant I-1823 and NIH grant MH61345 to SLS and the NIH grant 1S10OD021685 to the EM Core facility. We acknowledge Aparna Mohanakrishnan for the technical support in PCR and Heather Grossman for the technical support in adenovirus production.

## Author Contributions

ZC and SLS designed the experiments, interpreted the results and wrote the manuscript with input from all authors. ZC and MM performed the TIRF and WF microscope imaging and data analysis. ZC, RM and MB performed the TfnR/Tfn uptake and recycling assays. ZC performed the ITC, CCK-8 and pulldown assays. PM performed the ^125^I-LDL uptake assay. DKR performed the TEM imaging.

## Competing interests

The authors declare no competing interests.

## References

Aguet, F., C.N. Antonescu, M. Mettlen, S.L. Schmid, and G. Danuser. 2013. Advances in Analysis of Low Signal-to-Noise Images Link Dynamin and AP2 to the Functions of an Endocytic Checkpoint. Developmental Cell. 26:279–291.

Ahle, S., and E. Ungewickell. 1989. Identification of a clathrin binding subunit in the HA2 adaptor protein complex. Journal of Biological Chemistry. 264:20089–20093.

Bendris, N., K.C. Williams, C.R. Reis, E.S. Welf, P.H. Chen, B. Lemmers, M. Hahne, H.S. Leong, and S.L. Schmid. 2016. SNX9 promotes metastasis by enhancing cancer cell invasion via differential regulation of RhoGTPases. Mol Biol Cell.

Bennett, E.M., S.X. Lin, M.C. Towler, F.R. Maxfield, and F.M. Brodsky. 2001. Clathrin Hub Expression Affects Early Endosome Distribution with Minimal Impact on Receptor Sorting and Recycling. Molecular Biology of the Cell. 12:2790–2799.

Bitsikas, V., I.R. Corrêa, Jr., and B.J. Nichols. 2014. Clathrin-independent pathways do not contribute significantly to endocytic flux. eLife. 3:e03970.

Brodsky, F.M., C.-Y. Chen, C. Knuehl, M.C. Towler, and D.E. Wakeham. 2001. Biological Basket Weaving: Formation and Function of Clathrin-Coated Vesicles. Annu Rev Cell Dev Bi. 17:517–568.

Chen, P.-H., N. Bendris, Y.-J. Hsiao, C.R. Reis, M. Mettlen, H.-Y. Chen, S.-L. Yu, and S.L. Schmid. 2017. Crosstalk between CLCb/Dyn1-Mediated Adaptive Clathrin-Mediated Endocytosis and Epidermal Growth Factor Receptor Signaling Increases Metastasis. Developmental Cell. 40:278–288.e275.

Cocucci, E., F. Aguet, S. Boulant, and T. Kirchhausen. 2012. The First Five Seconds in the Life of a Clathrin-Coated Pit. Cell. 150:495–507.

Collette, J.R., R.J. Chi, D.R. Boettner, I.M. Fernandez-Golbano, R. Plemel, A.J. Merz, M.I. Geli, L.M. Traub, and S.K. Lemmon. 2009. Clathrin Functions in the Absence of the Terminal Domain Binding Site for Adaptor-associated Clathrin-Box Motifs. Molecular Biology of the Cell. 20:3401–3413.

Conner, S.D., and S.L. Schmid. 2003. Differential requirements for AP-2 in clathrin-mediated endocytosis. J Cell Biol. 162:773–779.

Copolovici, D.M., K. Langel, E. Eriste, and U. Langel. 2014. Cell-Penetrating Peptides: Design, Synthesis, and Applications. Acs Nano. 8:1972–1994.

Damke, H., T. Baba, A.M. van der Bliek, and S.L. Schmid. 1995. Clathrin-independent pinocytosis is induced in cells overexpressing a temperature-sensitive mutant of dynamin. The Journal of cell biology. 131:69–80.

Damke, H., D.D. Binns, H. Ueda, S.L. Schmid, and T. Baba. 2001. Dynamin GTPase domain mutants block endocytic vesicle formation at morphologically distinct stages. Mol Biol Cell. 12:2578–2589.

Dell’Angelica, E.C. 2001. Clathrin-binding proteins: Got a motif? Join the network! Trends in Cell Biology. 11:315–318.

DeMari, J., C. Mroske, S. Tang, J. Nimeh, R. Miller, and R.R. Lebel. 2016. CLTC as a clinically novel gene associated with multiple malformations and developmental delay. American Journal of Medical Genetics Part A. 170:958–966.

Dutta, D., C.D. Williamson, N.B. Cole, and J.G. Donaldson. 2012. Pitstop 2 Is a Potent Inhibitor of Clathrin-Independent Endocytosis. PLOS ONE. 7:e45799.

Edeling, M.A., S.K. Mishra, P.A. Keyel, A.L. Steinhauser, B.M. Collins, R. Roth, J.E. Heuser, D.J. Owen, and L.M. Traub. 2006. Molecular Switches Involving the AP-2 β2 Appendage Regulate Endocytic Cargo Selection and Clathrin Coat Assembly. Developmental Cell. 10:329–342.

Fotin, A., Y. Cheng, P. Sliz, N. Grigorieff, S.C. Harrison, T. Kirchhausen, and T. Walz. 2004. Molecular model for a complete clathrin lattice from electron cryomicroscopy. Nature. 432:573–579.

Gallusser, A., and T. Kirchhausen. 1993. The beta 1 and beta 2 subunits of the AP complexes are the clathrin coat assembly components. The EMBO journal. 12:5237–5244.

Godlee, C., and M. Kaksonen. 2013. From uncertain beginnings: Initiation mechanisms of clathrin-mediated endocytosis. The Journal of Cell Biology. 203:717–725.

Greene, B., S.-H. Liu, A. Wilde, and F.M. Brodsky. 2000. Complete Reconstitution of Clathrin Basket Formation with Recombinant Protein Fragments: Adaptor Control of Clathrin Self-Assembly. Traffic. 1:69–75.

Hamdan, F.F., C.T. Myers, P. Cossette, P. Lemay, D. Spiegelman, A.D. Laporte, C. Nassif, O. Diallo, J. Monlong, M. Cadieux-Dion, S. Dobrzeniecka, C. Meloche, K. Retterer, M.T. Cho, J.A. Rosenfeld, W. Bi, C. Massicotte, M. Miguet, L. Brunga, B.M. Regan, K. Mo, C. Tam, A. Schneider, G. Hollingsworth, D.R. FitzPatrick, A. Donaldson, N. Canham, E. Blair, B. Kerr, A.E. Fry, R.H. Thomas, J. Shelagh, J.A. Hurst, H. Brittain, M. Blyth, R.R. Lebel, E.H. Gerkes, L. Davis-Keppen, Q. Stein, W.K. Chung, S.J. Dorison, P.J. Benke, E. Fassi, N. Corsten-Janssen, E.-J. Kamsteeg, F.T. Mau-Them, A.-L. Bruel, A. Verloes, K. Õunap, M.H. Wojcik, D.V.F. Albert, S. Venkateswaran, T. Ware, D. Jones, Y.-C. Liu, S.S. Mohammad, P. Bizargity, C.A. Bacino, V. Leuzzi, S. Martinelli, B. Dallapiccola, M. Tartaglia, L. Blumkin, K.J. Wierenga, G. Purcarin, J.J. O’Byrne, S. Stockler, A. Lehman, B. Keren, M.-C. Nougues, C. Mignot, S. Auvin, C. Nava, S.M. Hiatt, M. Bebin, Y. Shao, F. Scaglia, S.R. Lalani, R.E. Frye, I.T. Jarjour, S. Jacques, R.-M. Boucher, E. Riou, M. Srour, L. Carmant, A. Lortie, P. Major, P. Diadori, F. Dubeau, G. D’Anjou, G. Bourque, S.F. Berkovic, L.G. Sadleir, P.M. Campeau, Z. Kibar, R.G. Lafrenière, S.L. Girard, S. Mercimek-Mahmutoglu, C. Boelman, G.A. Rouleau, et al. 2017. High Rate of Recurrent De Novo Mutations in Developmental and Epileptic Encephalopathies. The American Journal of Human Genetics. 101:664–685.

Jaqaman, K., D. Loerke, M. Mettlen, H. Kuwata, S. Grinstein, S.L. Schmid, and G. Danuser. 2008. Robust single-particle tracking in live-cell time-lapse sequences. Nature Methods. 5:695–702.

Kadlecova, Z., S.J. Spielman, D. Loerke, A. Mohanakrishnan, D.K. Reed, and S.L. Schmid. 2017. Regulation of clathrin-mediated endocytosis by hierarchical allosteric activation of AP2. J Cell Biol. 216:167–179.

Kang, D.S., R.C. Kern, M.A. Puthenveedu, M. von Zastrow, J.C. Williams, and J.L. Benovic. 2009. Structure of an Arrestin2-Clathrin Complex Reveals a Novel Clathrin Binding Domain That Modulates Receptor Trafficking. Journal of Biological Chemistry. 284:29860–29872.

Kelly, B.T., S.C. Graham, N. Liska, P.N. Dannhauser, S. Höning, E.J. Ungewickell, and D.J. Owen. 2014. AP2 controls clathrin polymerization with a membrane-activated switch. Science. 345:459–463.

Kelly, B.T., A.J. McCoy, K. Späte, S.E. Miller, P.R. Evans, S. Höning, and D.J. Owen. 2008. A structural explanation for the binding of endocytic dileucine motifs by the AP2 complex. Nature. 456:976.

Keyel, P.A., S.K. Mishra, R. Roth, J.E. Heuser, S.C. Watkins, and L.M. Traub. 2006. A Single Common Portal for Clathrin-mediated Endocytosis of Distinct Cargo Governed by Cargo-selective Adaptors. Molecular Biology of the Cell. 17:4300–4317.

Kirchhausen, T., and S.C. Harrison. 1981. Protein organization in clathrin trimers. Cell. 23:755–761.

Lemmon, S.K., and L.M. Traub. 2012. Getting in Touch with the Clathrin Terminal Domain. Traffic. 13:511–519.

Liashkovich, I., D. Pasrednik, V. Prystopiuk, G. Rosso, H. Oberleithner, and V. Shahin. 2015. Clathrin inhibitor Pitstop-2 disrupts the nuclear pore complex permeability barrier. Sci Rep-Uk. 5.

Liu, S.-H., M.S. Marks, and F.M. Brodsky. 1998. A Dominant-negative Clathrin Mutant Differentially Affects Trafficking of Molecules with Distinct Sorting Motifs in the Class II Major Histocompatibility Complex (MHC) Pathway. The Journal of Cell Biology. 140:1023–1037.

Loerke, D., M. Mettlen, S.L. Schmid, and G. Danuser. 2011. Measuring the hierarchy of molecular events during clathrin-mediated endocytosis. Traffic. 12:815–825.

Lombardi, P., M. Mulder, H. van der Boom, R.R. Frants, and L.M. Havekes. 1993. Inefficient degradation of triglyceride-rich lipoprotein by HepG2 cells is due to a retarded transport to the lysosomal compartment. Journal of Biological Chemistry. 268:26113–26119.

Mattera, R., M. Boehm, R. Chaudhuri, Y. Prabhu, and J.S. Bonifacino. 2011. Conservation and Diversification of Dileucine Signal Recognition by Adaptor Protein (AP) Complex Variants. Journal of Biological Chemistry. 286:2022–2030.

Maurer, M.E., and J.A. Cooper. 2006. The adaptor protein Dab2 sorts LDL receptors into coated pits independently of AP-2 and ARH. Journal of Cell Science. 119:4235–4246.

Mayor, S., R.G. Parton, and J.G. Donaldson. 2014. Clathrin-independent pathways of endocytosis. Csh Perspect Biol. 6:a016758.

Merrifield, C.J., and M. Kaksonen. 2014. Endocytic Accessory Factors and Regulation of Clathrin-Mediated Endocytosis. Csh Perspect Biol. 6.

Mettlen, M., P.-H. Chen, S. Srinivasan, G. Danuser, and S.L. Schmid. 2018. Regulation of Clathrin-Mediated Endocytosis. Annual Review of Biochemistry. 87:871–896.

Mettlen, M., and G. Danuser. 2014. Imaging and Modeling the Dynamics of Clathrin-Mediated Endocytosis. Csh Perspect Biol. 6.

Michaely, P., Z. Zhao, W.-P. Li, R. Garuti, L.J. Huang, H.H. Hobbs, and J.C. Cohen. 2007. Identification of a VLDL-induced, FDNPVY-independent internalization mechanism for the LDLR. The EMBO journal. 26:3273–3282.

Miele, A.E., P.J. Watson, P.R. Evans, L.M. Traub, and D.J. Owen. 2004. Two distinct interaction motifs in amphiphysin bind two independent sites on the clathrin terminal domain β-propeller. Nature Structural & Molecular Biology. 11:242–248.

Moskowitz, H.S., C.T. Yokoyama, and T.A. Ryan. 2005. Highly cooperative control of endocytosis by clathrin. Molecular biology of the cell. 16:1769–1776.

Motley, A., N.A. Bright, M.N.J. Seaman, and M.S. Robinson. 2003. Clathrin-mediated endocytosis in AP-2-depleted cells. Journal of Cell Biology. 162:909–918.

Musacchio, A., C.J. Smith, A.M. Roseman, S.C. Harrison, T. Kirchhausen, and B.M.F. Pearse. 1999. Functional Organization of Clathrin in Coats: Combining Electron Cryomicroscopy and X-Ray Crystallography. Molecular Cell. 3:761–770.

Ohno, H., M.-C. Fournier, G. Poy, and J.S. Bonifacino. 1996. Structural Determinants of Interaction of Tyrosine-based Sorting Signals with the Adaptor Medium Chains. Journal of Biological Chemistry. 271:29009–29015.

Owen, D.J., and P.R. Evans. 1998. A Structural Explanation for the Recognition of Tyrosine-Based Endocytotic Signals. Science. 282:1327–1332.

Owen, D.J., Y. Vallis, M.E.M. Noble, J.B. Hunter, T.R. Dafforn, P.R. Evans, and H.T. McMahon. 1999. A Structural Explanation for the Binding of Multiple Ligands by the α-Adaptin Appendage Domain. Cell. 97:805–815.

Owen, D.J., Y. Vallis, B.M. Pearse, H.T. McMahon, and P.R. Evans. 2000. The structure and function of the beta 2-adaptin appendage domain. The EMBO journal. 19:4216–4227.

Park, R.J., H. Shen, L. Liu, X. Liu, S.M. Ferguson, and P. De Camilli. 2013. Dynamin triple knockout cells reveal off target effects of commonly used dynamin inhibitors. Journal of Cell Science. 126:5305–5312.

Pearse, B.M. 1982. Coated vesicles from human placenta carry ferritin, transferrin, and immunoglobulin G. Proceedings of the National Academy of Sciences. 79:451–455.

Peden, A.A., R.E. Rudge, W.W.Y. Lui, and M.S. Robinson. 2002. Assembly and function of AP-3 complexes in cells expressing mutant subunits. The Journal of cell biology. 156:327–336.

Persaud, A., Y. Cormerais, J. Pouyssegur, and D. Rotin. 2018. Dynamin inhibitors block activation of mTORC1 by amino acids independently of dynamin. Journal of Cell Science. 131:jcs211755.

Praefcke, G.J.K., M.G.J. Ford, E.M. Schmid, L.E. Olesen, J.L. Gallop, S.-Y. Peak-Chew, Y. Vallis, M.M. Babu, I.G. Mills, and H.T. McMahon. 2004. Evolving nature of the AP2 alpha-appendage hub during clathrin-coated vesicle endocytosis. The EMBO journal. 23:4371–4383.

Rosselli-Murai, L.K., J.A. Yates, S. Yoshida, J. Bourg, K.K.Y. Ho, M. White, J. Prisby, X. Tan, M. Altemus, L. Bao, Z.-F. Wu, S.L. Veatch, J.A. Swanson, S.D. Merajver, and A.P. Liu. 2018. Loss of PTEN promotes formation of signaling-capable clathrin-coated pits. Journal of Cell Science. 131:jcs208926.

Royle, S.J. 2006. The cellular functions of clathrin. Cell Mol Life Sci. 63:1823–1832.

Schmid, E.M., M.G.J. Ford, A. Burtey, G.J.K. Praefcke, S.-Y. Peak-Chew, I.G. Mills, A. Benmerah, and H.T. McMahon. 2006. Role of the AP2 β-Appendage Hub in Recruiting Partners for Clathrin-Coated Vesicle Assembly. PLOS Biology. 4:e262.

Schmid, E.M., and H.T. McMahon. 2007. Integrating molecular and network biology to decode endocytosis. Nature. 448:883–888.

Schmid, S.L., and E. Smythe. 1991. Stage-specific assays for coated pit formation and coated vesicle budding in vitro. The Journal of Cell Biology. 114:869–880.

Schöneberg, J., M. Lehmann, A. Ullrich, Y. Posor, W.-T. Lo, G. Lichtner, J. Schmoranzer, V. Haucke, and F. Noé. 2017. Lipid-mediated PX-BAR domain recruitment couples local membrane constriction to endocytic vesicle fission. Nature Communications. 8:15873.

Shih, W., A. Gallusser, and T. Kirchhausen. 1995. A Clathrin-binding Site in the Hinge of the 2 Chain of Mammalian AP-2 Complexes. Journal of Biological Chemistry. 270:31083–31090.

Smith, C.M., V. Haucke, A. McCluskey, P.J. Robinson, and M. Chircop. 2013. Inhibition of clathrin by pitstop 2 activates the spindle assembly checkpoint and induces cell death in dividing HeLa cancer cells. Mol Cancer. 12.

Srinivasan, S., C.J. Burckhardt, M. Bhave, Z.M. Chen, P.H. Chen, X.X. Wang, G. Danuser, and S.L. Schmid. 2018. A noncanonical role for dynamin-1 in regulating early stages of clathrin-mediated endocytosis in non-neuronal cells. Plos Biology. 16.

Takahashi, E., A. Haga, and H. Tanihara. 2015. Merlin Regulates Epithelial-to-Mesenchymal Transition of ARPE-19 Cells via TAK1-p38MAPK-Mediated Activation. Investigative Ophthalmology & Visual Science. 56:2449–2458.

ter Haar, E., S.C. Harrison, and T. Kirchhausen. 2000. Peptide-in-groove interactions link target proteins to the β-propeller of clathrin. Proceedings of the National Academy of Sciences. 97:1096–1100.

Traub, L.M., and J.S. Bonifacino. 2013. Cargo Recognition in Clathrin-Mediated Endocytosis. Csh Perspect Biol. 5.

Traub, L.M., M.A. Downs, J.L. Westrich, and D.H. Fremont. 1999. Crystal structure of the alpha appendage of AP-2 reveals a recruitment platform for clathrin-coat assembly. P Natl Acad Sci USA. 96:8907–8912.

Ungewickell, E., and D. Branton. 1981. Assembly units of clathrin coats. Nature. 289:420–422.

von Kleist, L., W. Stahlschmidt, H. Bulut, K. Gromova, D. Puchkov, M.J. Robertson, K.A. MacGregor, N. Tomilin, A. Pechstein, N. Chau, M. Chircop, J. Sakoff, J.P. von Kries, W. Saenger, H.G. Krausslich, O. Shupliakov, P.J. Robinson, A. McCluskey, and V. Haucke. 2011. Role of the Clathrin Terminal Domain in Regulating Coated Pit Dynamics Revealed by Small Molecule Inhibition (vol 146, pg 471, 2011). Cell. 146:841–841.

Walrant, A., S. Cardon, F. Burlina, and S. Sagan. 2017. Membrane Crossing and Membranotropic Activity of Cell-Penetrating Peptides: Dangerous Liaisons? Accounts Chem Res. 50:2968–2975.

Willox, A.K., and S.J. Royle. 2012. Functional Analysis of Interaction Sites on the N-Terminal Domain of Clathrin Heavy Chain. Traffic. 13:70–81.

Willox, A.K., Y.M.E. Sahraoui, and S.J. Royle. 2014. Non-specificity of Pitstop 2 in clathrin-mediated endocytosis. Biol Open. 3:326–331.

Wood, L.A., G. Larocque, N.I. Clarke, S. Sarkar, and S.J. Royle. 2017. New tools for “hot-wiring” clathrin-mediated endocytosis with temporal and spatial precision. The Journal of Cell Biology. 216:4351–4365.

Yarar, D., C.M. Waterman-Storer, and S.L. Schmid. 2007. SNX9 couples actin assembly to phosphoinositide signals and is required for membrane remodeling during endocytosis. Dev Cell. 13:43–56.

Zhuo, Y., U. Ilangovan, V. Schirf, B. Demeler, R. Sousa, A.P. Hinck, and E.M. Lafer. 2010. Dynamic Interactions between Clathrin and Locally Structured Elements in a Disordered Protein Mediate Clathrin Lattice Assembly. Journal of Molecular Biology. 404:274–290.

