## Supplemental Materials for "Structure-based inhibitors reveal roles for the clathrin terminal domain and its W-box binding site in CME"

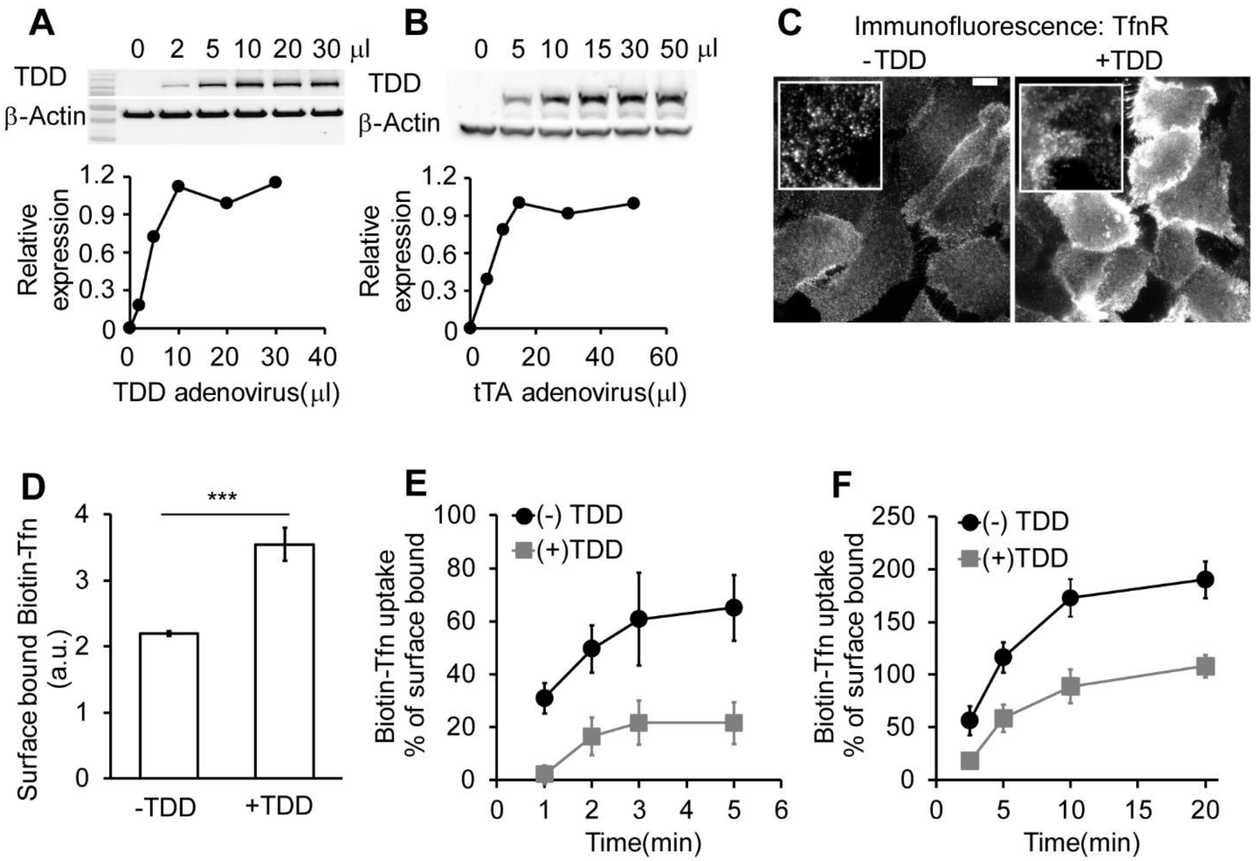


**Figure S1. Optimization of recombinant adeno-viral system in ARPE cells.** The amounts of TDD and helper tTA adeno-viruses were optimized for uniform infection of ARPE cells. (A) Relative TDD expression level after infection with 12μl tTA recombinant adenovirus and the indicated volumes of TDD recombinant adenoviruses. (B) Relative TDD expression level after infection with 20μl TDD recombinant adenovirus and the indicated volumes of tTA recombinant adenoviruses. (C) Immunofluorescence of surface bound TfnR with or without TDD expression. (D-F) Effects of TDD overexpression on Biotin-Tfn uptake revealed that TDD expression enhanced surface bound Biotin-Tfn (D) and reduced its uptake efficiency in single-round (E) and multi-round (F) Biotin-Tfn uptake. Average value ± standard deviations are from N=4 replicates.

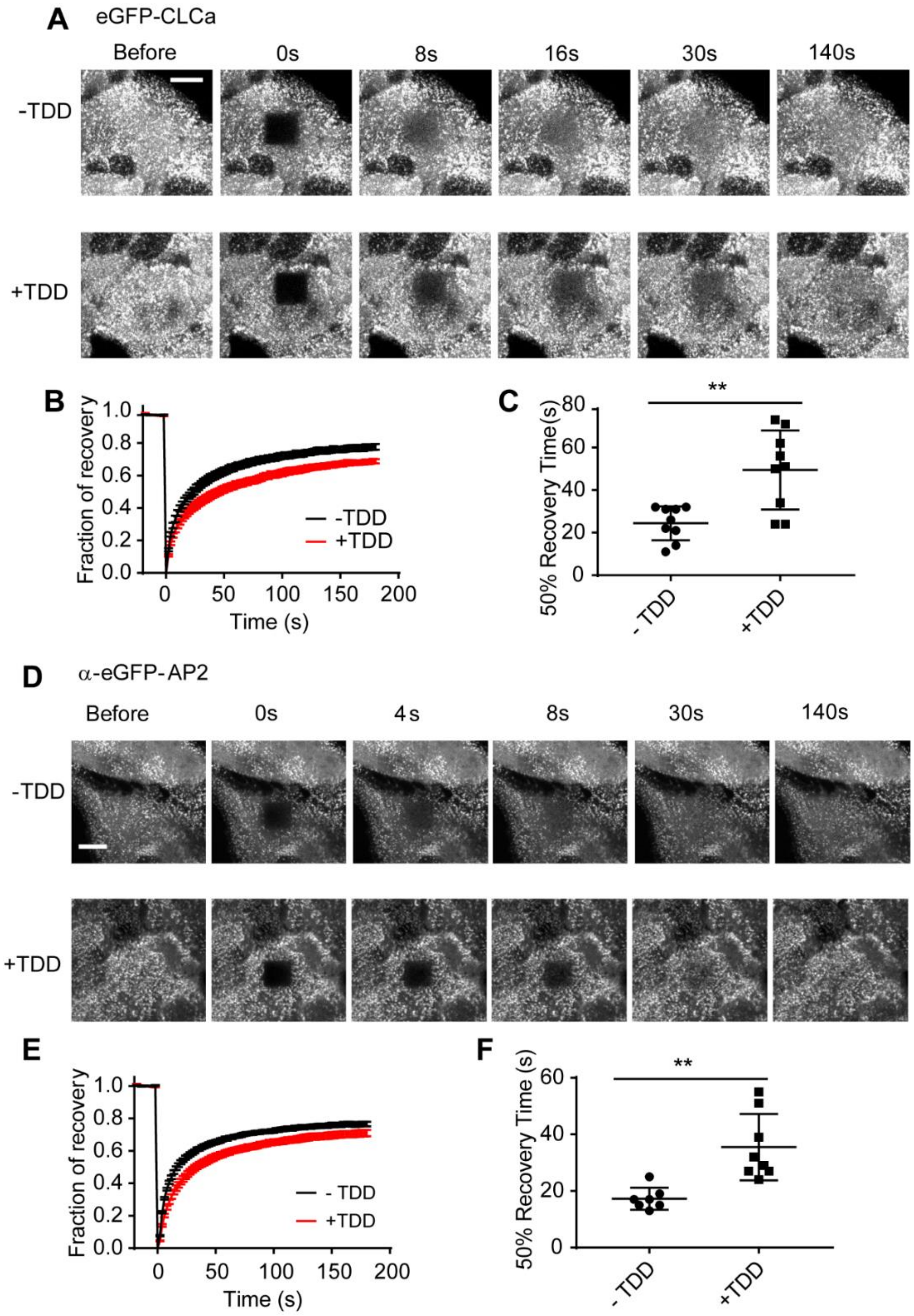


**Figure S2**. **Clathrin and AP2 exchange on plasma membrane is inhibited by TDD expression**. (A-C) Fluorescence Recovery After Photobleaching (FRAP) was conducted at 37°C in ARPE/HPV eGFP-CLCa cells with or without TDD overexpression using confocal microscope. (A) Representative time-lapse FRAP images. (B) Average fluorescent intensity traces of the photobleached area (dark square at t = 0s). (C) Time required for 50% fluorescence recovery. (D-F) FRAP was conducted at 37°C in ARPE α-eGFP-AP2 cells with or without TDD overexpression using confocal microscope. (D) Representative time-lapse FRAP images. (E) Average fluorescent intensity traces of the photobleached area (dark square at t = 0s). (F) Time required for 50% fluorescence recovery. Error bars are standard deviations from N=8 experiments. Scale bars =3μm.

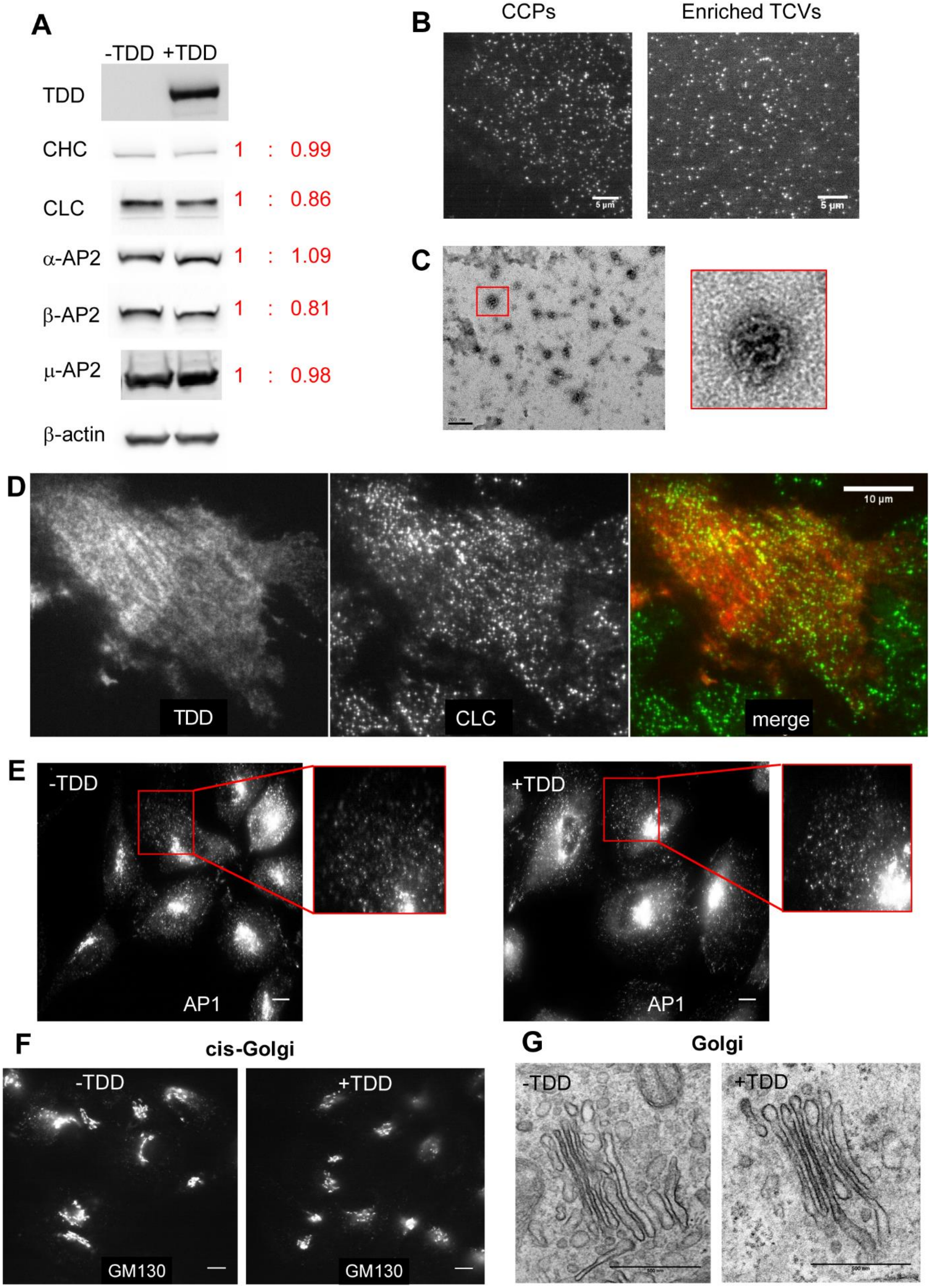
A
-TDD +TDD
TDD
CHC 1 : 0.99
CLC 1 : 0.86
α-AP2 1 : 1.09
β-AP2 1 : 0.81
μ-AP2 1 : 0.98
β-actin
B
CCPs
Enriched TCVs
5 μm
5 μm
C
D
TDD
CLC
merge
10 μm
E
-TDD
AP1
+TDD
AP1
F
cis-Golgi
-TDD
GM130
+TDD
GM130
G
Golgi
-TDD
+TDD

**Figure S3. Effect of TDD expression on endogenous protein expression, clathrin-coat stability, and AP1-mediated clathrin trafficking from Golgi.** (A) ARPE/HPV cells expressing eGFP-CLCa cells were treated with or without TDD overexpression. TDD expression did not alter the expression levels of endogenous clathrin or AP2. (B) Representative TIRF images of CCPs in live cells or isolated TCVs. Scale bars = 5µm. (C) Representative electron microscopy images showing the collapsed coats typical of isolated TCVs. Scale bars = 200nm. (D) Immunostaining of HA-TDD in ARPE/HPV eGFP-CLCa cells. Scale bar = 10µm. (E-G) ARPE cells were treated with or without TDD overexpression. Immunostaining of AP1 γ subunit (E) and GM-130 cis-Golgi marker (F) were imaged with wide field microscopy. Scale bars = 10µm. (G) Representative electron microscopy images of Golgi apparatus. Scale bars = 500nm.

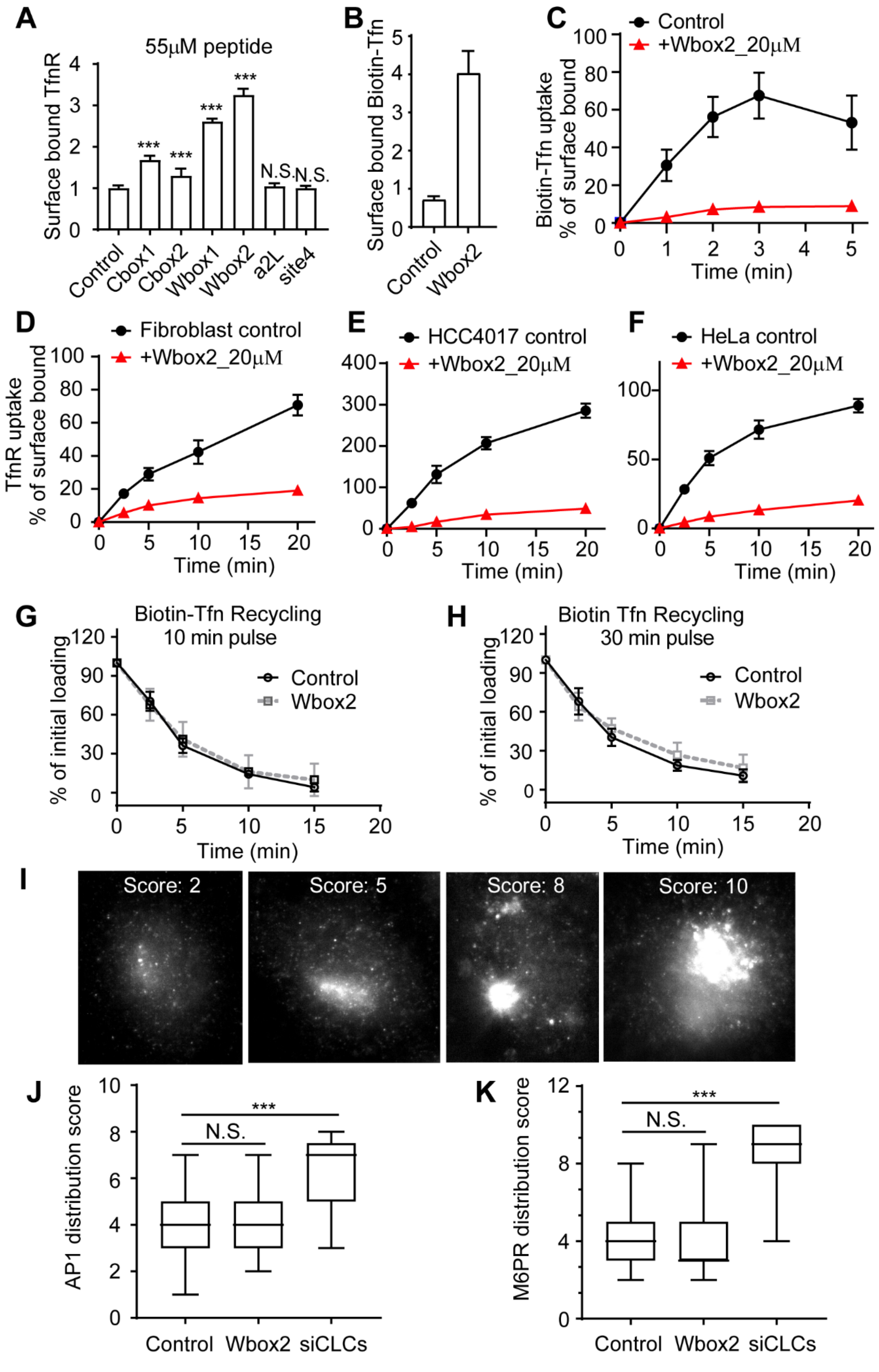

A
55μM peptide
Surface bound TfnR
***
***
***
***
N.S.
N.S.
Control
Cbox1
Cbox2
Wbox1
Wbox2
a2L
site4
B
Surface bound Biotin-Tfn
Control
Wbox2
C
Control
+Wbox2_20μM
Biotin-Tfn uptake
% of surface bound
Time (min)
D
Fibroblast control
+Wbox2_20μM
TfnR uptake
% of surface bound
Time (min)
E
HCC4017 control
+Wbox2_20μM
Time (min)
F
HeLa control
+Wbox2_20μM
Time (min)
G
Biotin-Tfn Recycling
10 min pulse
% of initial loading
Control
Wbox2
Time (min)
H
Biotin Tfn Recycling
30 min pulse
% of initial loading
Control
Wbox2
Time (min)
I
Score: 2
Score: 5
Score: 8
Score: 10
J
AP1 distribution score
***
N.S.
Control
Wbox2
siCLCs
K
M6PR distribution score
***
N.S.
Control
Wbox2
siCLCs

**Figure S4. Effect of TD-derived peptides on CME, Tfn recycling and AP1-mediated Golgi trafficking.** (A) Effect of the inhibitory activities of the 6 peptides on the surface bound TfnR. (B-C) Single round Biotin-Tfn uptake revealed that Wbox2 strongly enhanced surface bound Biotin-Tfn (B) and reduced Biotin-Tfn uptake efficiency (C). (D-F) The Wbox2 peptide inhibited TfnR uptake in Fibroblast, HCC4017 and HeLa cells. (G-H) Effects of Wbox2 on Biotin-Transferrin recycling after either a 10 min and 30min pulse. (I-K) Immunostaining of AP1 γ-adaptin and Mannose-6-phosphate-receptor (M6PR) in control HeLa cells as well as cells treated with Wbox2 (10μM, pre-incubation for 30min at 37°C) or siRNA knockdown CLCa+b (siCLCs). The AP1 and M6RP distribution in cells was quantified by grading them on the basis of degree of concentration at perinuclear region. Lowest score for completely dispersed phenotype, highest score for majority of signal being concentrated at perinuclear region, as representative images of M6PR shown in (I). (J-K) Wbox2 treatment did not alter AP1 or M6PR distribution, while siCLCa+b as a positive control showed a strong effect by accumulating AP1 and M6PR in the perinuclear region. Number of cells quantified in (J): 94 for Control, 106 for Wbox2, and 33 for siCLCs. Number of cells quantified in (K): 95 for Control, 62 for Wbox2, and 37 for siCLCs.

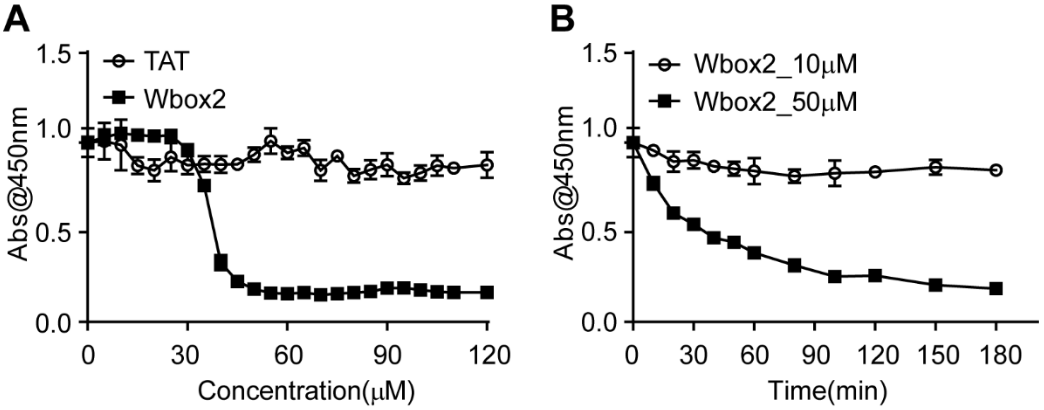


**Figure S5. Cell viability after Wbox2 treatment.** (A) Quantitation of viable cell number after treatment with varied concentrations of TAT and Wbox2 peptides using Cell Counting Kit 8 (CCK-8). (B) Quantification of viable cell number after treatment with 10 or 50uM of Wbox2 and varied incubation times. Average data ± standard deviations are from N=4 replicates.

**Supplemental Tables**

**Table S1** List of proteins involved in clathrin-mediated endocytosis pathway from three independent trials.

**Table S2** List of proteins involved in membrane trafficking pathways from three independent trials.

**Table S3** List of reagents used in this work, including chemicals, proteins, antibodies and siRNAs.

**Supplemental Videos**

**Video S1** Time-lapse TIRFM imaging of stable ARPE/HPV cells expressing eGFP-CLCa under control conditions. Cells were infected with TDD and tTA-encoding adenviruses but incubated in the presence of 100 ng/ml tetracycline to suppress TDD expression. Images were obtained at 1 frame/sec and collected for 7.5 min. Video is accelerated 50-fold.

**Video S2** Time-lapse TIRFM imaging of stable ARPE/HPV cells expressing eGFP-CLC. Cells were infected with TDD and tTA-encoding adenviruses but incubated in the absence of tetracycline to induce TDD expression. Images were obtained at 1 frame/sec and collected for 7.5 min. Video is accelerated 50-fold.

**Video S3** Time-lapse TIRFM imaging of stable ARPE cells expressing α-eGFP-AP2 under control conditions. Cells were infected with TDD and tTA-encoding adenviruses but incubated in the presence of 100 ng/ml tetracycline to suppress TDD expression. Images were obtained at 1 frame/sec and collected for 7.5 min. Video is accelerated 50-fold.

**Video S4** Time-lapse TIRFM imaging of stable ARPE cells expressing α-eGFP-AP2. Cells were infected with TDD and tTA-encoding adenviruses but incubated in the absence of tetracycline to induce TDD expression. Images were obtained at 1 frame/sec and collected for 7.5 min. Video is accelerated 50-fold.

**Video S5** Time-lapse TIRFM imaging of stable ARPE/HPV cells expressing eGFP-CLC 5 days after siRNA-mediated knockdown of SNX9. Images were obtained at 1 frame/sec and collected for 7.5 min. Video is accelerated 50-fold.

**Video S6** Time-lapse TIRFM imaging of stable in ARPE cells expressing α-eGFP-AP2 and mRuby-CLCa incubated in the presence of 20 µM CMEpi. Left shows α-eGFP-AP2 channel and right is mRuby-CLCa channel. Images were obtained at 1 frame/sec and collected for 7.5 min. Video is accelerated 50-fold.
